# Resistance diagnostics as a public health tool to combat antibiotic resistance: A model-based evaluation

**DOI:** 10.1101/452656

**Authors:** David McAdams, Kristofer Wollein Waldetoft, Christine Tedijanto, Marc Lipsitch, Sam P. Brown

## Abstract

Rapid point-of-care resistance diagnostics (POC-RD) are a key tool in the fight against antibiotic resistance. By tailoring drug choice to infection genotype, doctors can improve treatment efficacy while limiting costs of inappropriate antibiotic prescription. Here we combine epidemiological theory and data to assess the potential of RD innovations in a public health context, as a means to limit or even reverse selection for antibiotic resistance. POC-RD can be used to impose a non-biological fitness cost on resistant strains, by enabling diagnostic-informed treatment and targeted interventions that reduce resistant strains’ opportunities for transmission. We assess this diagnostic-imposed fitness cost in the context of a spectrum of bacterial population biologies, and find that the expected impact varies from selection against resistance for obligate pathogens to marginal public health improvements for opportunistic pathogens with high ‘bystander’ antibiotic exposure during asymptomatic carriage (e.g. the pneumococcus). We close by generalizing the notion of RD-informed strategies to incorporate carriage surveillance information, and illustrate that coupling transmission-control interventions to the discovery of resistant strains in carriage can potentially select against resistance in a broad range of opportunistic pathogens.

## Introduction

> “Because antibiotic resistance occurs as part of a natural evolution process, it can be significantly slowed but not stopped. Therefore, new antibiotics will always be needed to keep up with resistant bacteria.” (CDC, “Antibiotic Resistance Threats in the United States, 2013”) (1)

The antimicrobial resistance crisis threatens to undermine key features of modern medicine, at great costs in terms of patient morbidity, mortality and treatment expense (2–6). Current mainstream antibiotic-treatment strategies sow the seeds of their own downfall by strongly selecting for resistant strains, leading some to argue that continual new-antibiotic discovery is the only way to stay ahead of a “post-antibiotic future” (1, 7, 8). If this bleak vision is correct, there is an urgent need to buy time by extending the lifespan of existing antibiotics while research and development for new ones takes its course. More optimistically, it may be possible to improve how we use existing antibiotics and to implement other control measures so that an endless supply of new antibiotics is not required.

Among a number of innovative approaches to improve antibiotic use (9–15), one of the most promising leverages point-of-care resistance diagnostics (POC-RD) that provide prescribers with a rapid readout of the sensitivity/resistance profile of an infecting organism. POC-RD allows prescribers to choose older, cheaper, and/or narrower-spectrum antibiotics when such drugs are most appropriate for patients, thereby saving newer, more expensive, and/or broader-spectrum antibiotics for situations where they are really needed, and perhaps reducing the intensity of selection for resistance to these drugs (16–18).

Less often considered is a second potential benefit of resistance diagnosis: to enable “Search & Destroy” (S&D) tactics to combat the most dangerous resistant pathogen strains, such as methicillin-resistant *Staphyloccus aureus* (MRSA) and carbapenem-resistant *Enterobacteriaceae* (CRE) (19–24). S&D strategies aim to identify and then isolate patients who are carrying problematic resistant strains until pathogen clearance can be confirmed. If resistant strains can be rapidly, accurately identified and their transmission curtailed by targeted infection-control measures, then S&D can create a *non-biological diagnostic-imposed fitness cost* borne only by targeted resistant strains. However, the magnitude of this fitness cost is hotly debated, especially in the context of MRSA control (25, 26), and, in any event, intensive medical interventions such as patient isolation are not a practical or economical option in many circumstances.

In this paper, we ask: when it is possible to create *net selection against resistance*, even when (i) there are no biological fitness costs associated with resistance, (ii) the best available treatment cannot be withheld from any patient, and (iii) all non-antibiotic intervention options are only moderately effective. We define “net selection against resistance” as maintaining the fitness of one or more resistant strains below that of the drug-sensitive strain, so that the frequency of these resistant strains will decline toward zero. This can be accomplished, generally speaking, when diagnostics permit medical personnel to artificially shape the pathogen fitness landscape so that resistant strains are disproportionately disadvantaged. We show that the potential to reduce or even reverse selection on resistance depends on two key factors. (i) <u>Pathogen lifestyle</u>: is symptomatic disease tightly coupled to transmission (obligate pathogen), or can the pathogen also transmit from an asymptomatic carriage phase (opportunistic pathogen)? (ii) <u>Pan resistance</u>: are untreatable pan-resistant strains already in circulation? Table 1 offers a preview of our main findings in a simple setting with two equally-effective antibiotic-treatment options, “drug 1” (first-line treatment to which resistance has already emerged in the target-pathogen population) and “drug 2” (second-line treatment to which resistance may or may not have already emerged).

**Table 1:** Preview of main findings.

| Lifestyle | Pan-resistant | Key findings |
| --- | --- | --- |
| Simple Obligate | No | Net selection against drug-1 resistance is **possible**. |
| Simple Obligate | Yes | Net selection against pan resistance is **possible only if** *either* there are substantial biological fitness costs associated with pan resistance *or* a highly-effective infection intervention (“isolation”) is available. |
| Opportunistic | No/Yes | Net selection against drug-1 resistance **may be impossible**, even if all those with sensitive infection are left untreated and all those with drug-1-resistant infection could be targeted for isolation—**unless interventions are also conditioned on asymptomatic carriage RD** |

**Table 2:** Notation.

| Notation | Details |
| --- | --- |
| *Pathogen strains and pathogen-lifestyle parameters* |  |
| $x$ | Drugs $x = 1, 2$ are available to treat infections caused by the target pathogen. |
| $X$ | Each strain is named for the drugs (if any) $X = 0, 1, 2, 12$ to which it is resistant: strain 0 is susceptible to both drugs (“pan-sensitive”); strain 1 is resistant only to drug 1; strain 2 is resistant only to drug 2; and strain 12 is |
|  | resistant to both drugs (“pan-resistant”). |
| $\beta^c, \beta^I > 0$ | Transmission rates during carriage and during infection, if uncontrolled and if no biological fitness costs |
| $f_X \geq 0$ | Biological fitness cost associated with strain $X$ resistance, reducing transmission during carriage and infection by a factor of $1 - f_X$ |
| $\gamma^c > 0$ | Baseline carriage clearance rate |
| $\phi_x^c > 0$, | Carriage clearance rate of drug-$x$-sensitive strain due to bystander exposure to drug $x$ |
| $d > 0$ | Rate at which infection develops from carriage (in the limit $d \rightarrow \infty$, we recover the SIS case) |
| $\gamma_0^I > 0$ | Baseline infection clearance rate when untreated or ineffectively treated |
| *Diagnosis, treatment, and transmission-control parameters* |  |
| $D \geq 0$ | Diagnostic delay, i.e., time from sample collection to result |
| $r_X \geq 0$ | Rate at which resistant strain $X$ is discovered while in the carriage state |
| $\gamma_x^I > \gamma_0^I$ | Infection clearance rate of drug-$x$-sensitive strain when treated with drug $x$ |
| $Z_X^I, Z_X^C \leq 1$ | Proportional reduction in transmission during infection ($Z_X^I$) or during asymptomatic carriage ($Z_X^C$) due to heightened transmission control (HTC$_X$) measures targeted against strain $X$ |
| *Key derived variables* |  |
| $R_{0,X}$ | Basic reproduction number of strain $X$ |
| $f_X^*$ | Threshold biological fitness cost for strain $X$ not to enjoy a reproductive advantage, i.e., $R_{0,X} > R_{0,0}$ if $f_X < f_X^*$ and $R_{0,X} < R_{0,0}$ if $f_X > f_X^*$ |

## Methods

We describe a mathematical model for a single pathogen species (“target pathogen”) with multiple strains having different antibiotic susceptibilities, in which health care providers and public health officials (hereafter “providers”) can shape the pathogen fitness landscape by tailoring treatment and transmission-control measures informed by resistance diagnosis (RD).

### Pathogen strains

Two antibiotics are available to treat infections caused by the target pathogen: drug 1 (first-line treatment that would be prescribed to all patients in the absence of resistance diagnosis) and drug 2 (second-line treatment). Resistance to drug 1 and potentially also to drug 2 has already emerged in the target-pathogen population but not yet reached fixation. In particular, there are up to four resistance profiles in circulation: an untreatable “pan-resistant” strain (strain 12); a “drug-1-resistant” strain that remains sensitive to drug 2 (strain 1); a “drug-2-resistant” strain that remains sensitive to drug 1 (strain 2); and a “pan-sensitive” strain that can be effectively treated with either drug (strain 0) (Fig 1A). The model encompasses a spectrum of pathogen lifestyles, from “simple obligate pathogens” (causing disease immediately after colonization, Fig 1A,B) to “opportunistic pathogens” (transmitting from an asymptomatic carriage phase as well as during symptomatic disease, Fig 1C).

**Figure 1.**
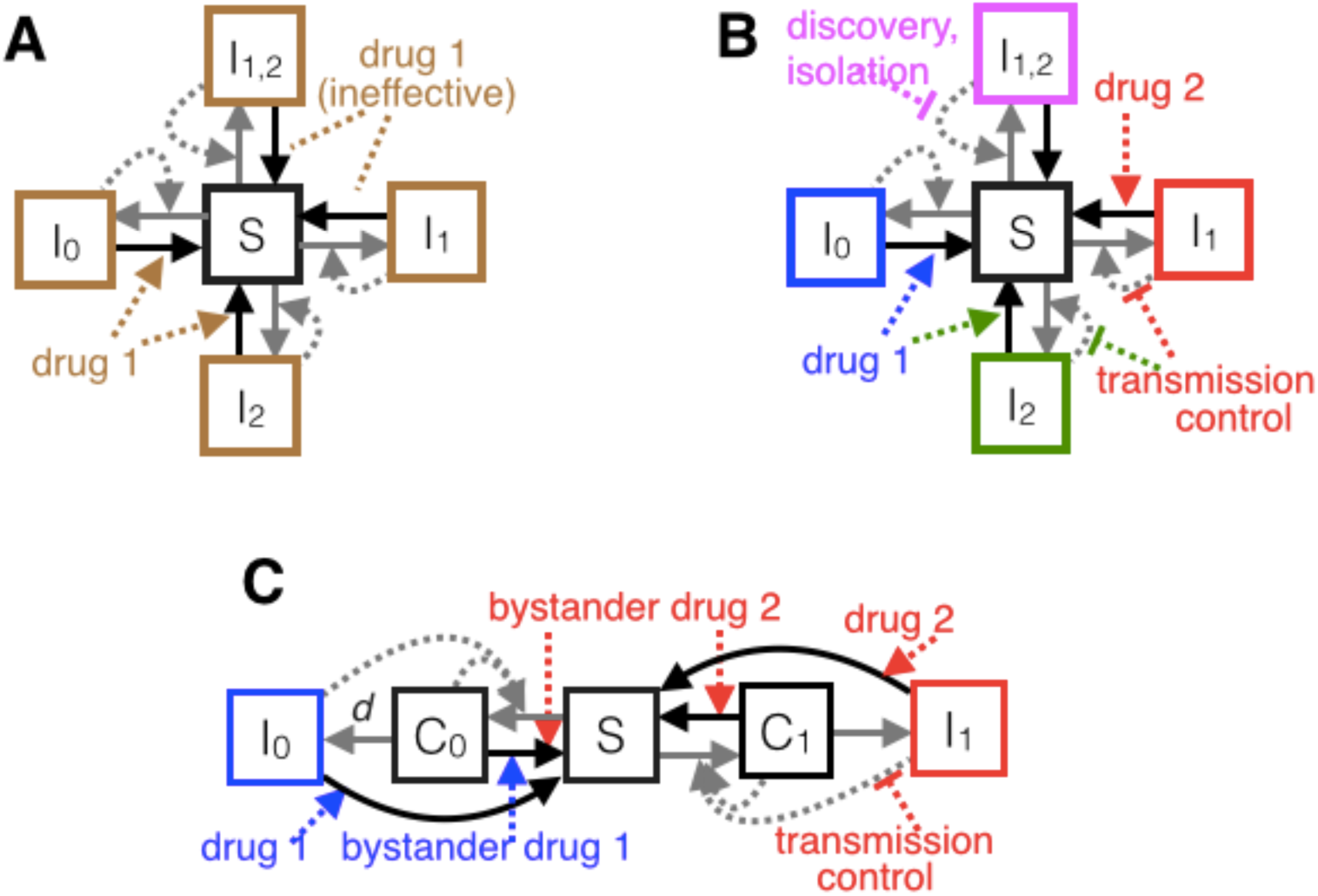
Schematic of the obligate/SIS (A,B) and opportunistic/SCIS (C) epidemiological models. Boxes denote proportions of hosts in mutually exclusive states: *S* for uninfected (susceptible) hosts, *I*_0_ for hosts infected with a strain sensitive to both drugs, and *I*_1_, *I*_2_, *I*_12_ for hosts infected with strains resistant to drugs 1, 2, or both 1 and 2, respectively. In the SCIS model (**C**, showing only two pathogen genotypes for clarity), *C*_*0*_ and *C*_1_ denote asymptomatic carriage of sensitive and drug-1-resistant bacteria, respectively, and *d* is the rate at which disease develops from carriage (when *d* → ∞, we recover an SIS model). Box colors denote distinct clinical presentations in the absence (**A**) or presence (**B,C**) of multi-drug POC-RD. Solid arrows represent flows of individuals between states, and dashed arrows represent factors influencing those flows (e.g., antibiotic treatment). Grey and black arrows denote transmission and clearance, respectively. Equations describing the system are in SI.B.

### Resistance diagnosis (RD)-informed treatment and control

We use standard reproduction-number analysis to investigate the pathogen-fitness impact of *RD-informed treatment and control*, depending on pathogen lifestyle (obligate vs opportunistic), what sort of resistance diagnostic is available, and what sort of transmission-control options can be feasibly targeted against each resistant strain, once identified. Cases considered include: point-of-care RD (POC-RD) with an obligate pathogen (Case #1, Fig 1B); POC-RD with an opportunistic pathogen (Case #2, Fig 1C); and carriage resistance surveillance (“carriage RD”) with an opportunistic pathogen (Case #3). See the SI for omitted mathematical details as well as several extensions, including to settings with intermediate resistance, public health interventions aimed at discovering drug-resistant infections more quickly, resistance-conferring mutation, host migration, and diagnostic escape.

## Results

### Case #1: POC-RD and a simple obligate pathogen (SIS model)

In the limiting case when the target pathogen immediately causes disease (*d* = ∞), what we call a “simple obligate pathogen,” our SCIS model reduces to a standard Susceptible-Infected (SIS) epidemiological model (27–29). Few if any real-world bacterial pathogens fit this case, but it is useful conceptually as a limiting case (Fig 1A,B).

Fig 2 illustrates the impact of RD-informed treatment and transmission control on whether the drug-1-resistant and/or pan-resistant strains enjoy a reproductive advantage relative to the pan-sensitive strain, for a generic pathogen with average duration of infection of 5 days (with effective treatment) or 10 days (without treatment). The blue parameter regions are where the drug-susceptible strain has a higher reproduction number (*R*_0_) than the drug-1-resistant strain and/or the pan-resistant strain, as a function of diagnostic delay *D* and the biological fitness costs *f*_1_, *f*_12_ of drug-1 and pan resistance, thereby creating net selection against these resistant strains.

**Figure 2.**
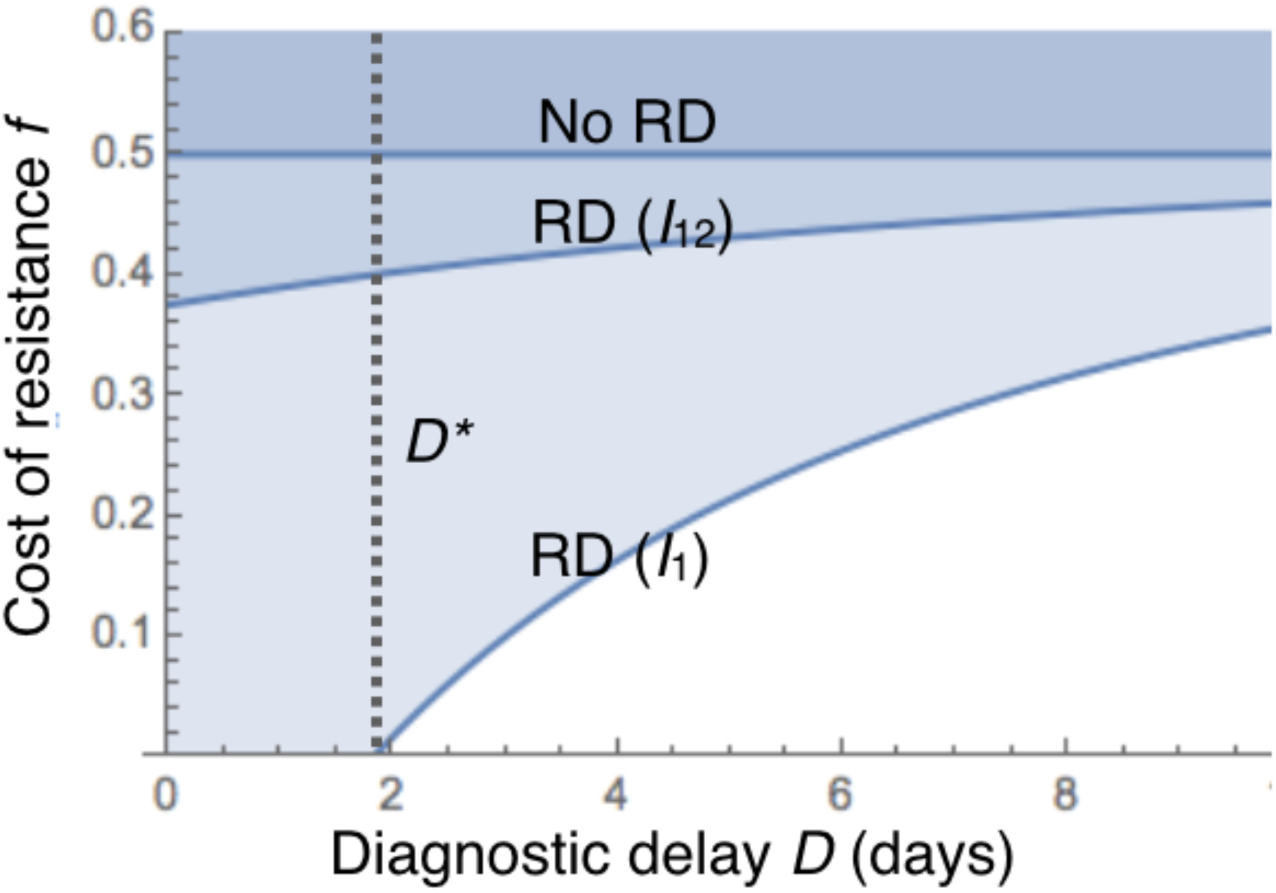
Rapid resistance diagnostics enable conditional treatment and infection control strategies that can select against resistance for obligate pathogens even with no biological costs of resistance. The minimal cost of resistance *f*\*(*D*) that allows universal treatment without causing an increase in resistance is plotted (contour lines) against diagnostic delay *D*. The dashed vertical line indicates the longest diagnostic delay (*D*^*^) given which there is selection against drug-1 resistance while treating all cases. Three scenarios are shown: resistance diagnostics (RD) not available (No RD, contour plot of 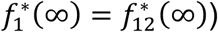; RD control of *I*_1_ only (RD (*I*_1_), contour plot of 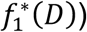; RD control of *I*_12_ (and trivially, *I*_1_) (RD (*I*_12_), contour plot of 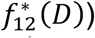. *R*_0_ expressions are defined in Box 1, with further details in SI.B. Parameters (rates per day): 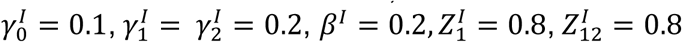.

**Box 1. Reproductive numbers (***<u>R</u>*<u>_0_</u>**) for strains in the SIS model.** See SI.B for more <u>detailed derivations</u>

##### Pan-sensitive strain

Strain-0 infections are treated with drug 1 and subjected to standard transmission control, whether or not RD is available. Since strain-0 infections clear at rate 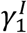 under drug-1 treatment and strain 0 transmits at rate *β*^*I*^ under standard control, strain 0’s reproduction number is

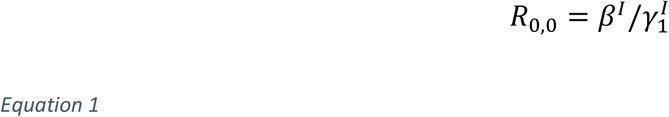

##### Drug-1-resistant strain

Since strain-1 infections clear at rate 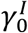 during the diagnostic-delay period, a patient will remain infected long enough for RD results to become available (triggering drug 2 and transmission control) with probability 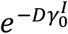. Depending on diagnostic delay *D* and fitness cost *f*_1_, strain 1’s reproduction number takes the form

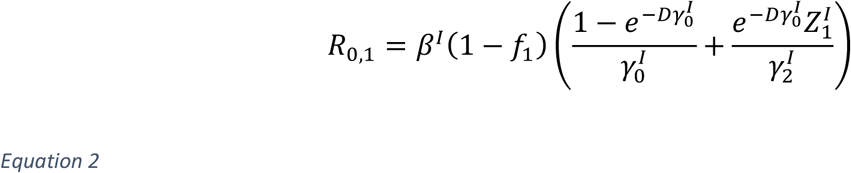

##### Drug-2-resistant strain

Since strain-2 infections are susceptible to drug 1 and therefore clear at rate 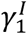 during the diagnostic-delay period, such infections will be diagnosed before clearance with probability 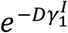. Strain 2’s resulting reproduction number is

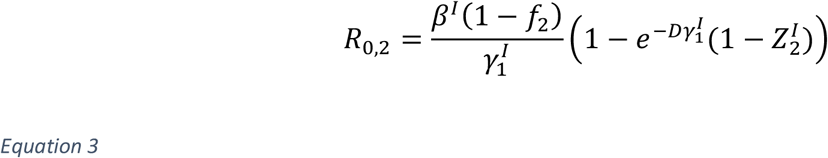

##### Pan-resistant strain

Strain-12 infections clear at the untreated rate 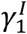 during and after the diagnostic-delay period. However, diagnostic results (if available before infection clearance, probability 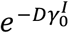) can trigger heightened transmission control, reducing strain 12’s reproduction number to

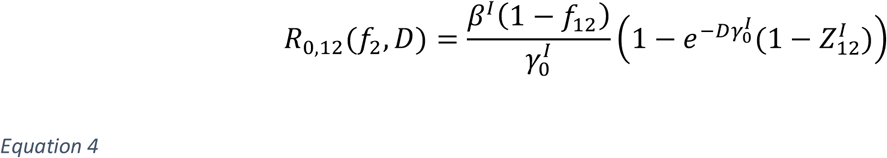

The solid contours in Fig 2 illustrate the critical threshold resistance costs for the drug-1-resistant strain 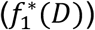 and the pan-resistant strain 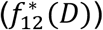, the smallest fitness costs that allow universal treatment while keeping the resistant strain at a reproductive disadvantage relative to the sensitive strain. The introduction of RD has the greatest impact on the drug 1 resistant strain (Fig 2, strain *I*1). In the limit of no diagnostic delay (*D* = 0), we find that 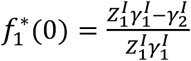. This implies that resistance to drug 1 (given available drug 2) can be selected against even in the absence of resistance costs 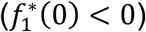 so long as *either* drug 2 is more effective than drug 1 *or* drug 2 is less effective than drug 1 but transmission control is sufficiently effective that 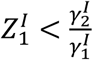.

Control of a circulating pan-resistant strain *I*12 is more challenging, even with POC-RD. Absent fitness costs (*f*_12_ = 0), net selection against pan resistance can be maintained only if 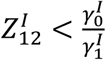, meaning that transmission control of the pan-resistant strain is more effective than drug-1 treatment at speeding clearance. In the case of more modest transmission control (Fig 2, 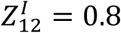), net selection against pan resistance requires that there be sufficient fitness costs associated with pan resistance. (In the POC-RD limit when *D* = 0, the threshold fitness cost 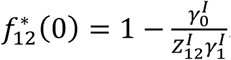)

Both thresholds 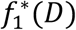 and 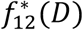 are increasing in *D*, as longer delays reduce the effectiveness of diagnostic-informed treatment and control, and converge in the *D* → ∞ limit to the “no-RD fitness-cost threshold” absent any resistance diagnosis: 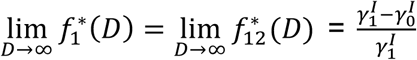. In Fig 2 with baseline clearance 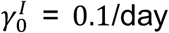 and drug 1 assisted clearance 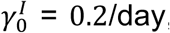, the no-RD fitness-cost threshold equals 0.5, greatly exceeding typical reported costs of resistance. To give some context on empirical estimates of costs of resistance, *f*, a recent meta-analysis estimated *f* = 0.21 (+/-0.024) for chromosomally encoded resistance and *f* = 0.09 (+/-0.024) for plasmid-encoded resistance (30), albeit using growth rate rather than epidemiological transmission measures of fitness effects.

### Case #2: an opportunistic pathogen with a carrier state (SCIS model)

Many disease-causing bacteria are opportunistic pathogens capable of transmission from an asymptomatic carriage state, while living harmlessly in a host microbial compartment such as the gut or the nasopharynx. Such pathogens face “bystander exposure” to antibiotics used to treat infections caused by other pathogens or to treat noninfectious conditions (31). Take for example the pneumococcus (*S. pneumoniae*), one of the top bacterial causes of death globally (32) and a leading cause of antibiotic prescription. Despite the severe burden of disease, the pneumococcus is subjected in the US to an estimated 9.1 times more courses of any antibiotic during asymptomatic carriage than during disease. Fig 3 compares the volume of bystander selection to target antibiotic exposure for several major bacterial pathogens. An alternate visualization of the proportions of bystander exposure (absent *C. difficile*) is presented in Tedijanto et al. (31).

**Figure 3.**
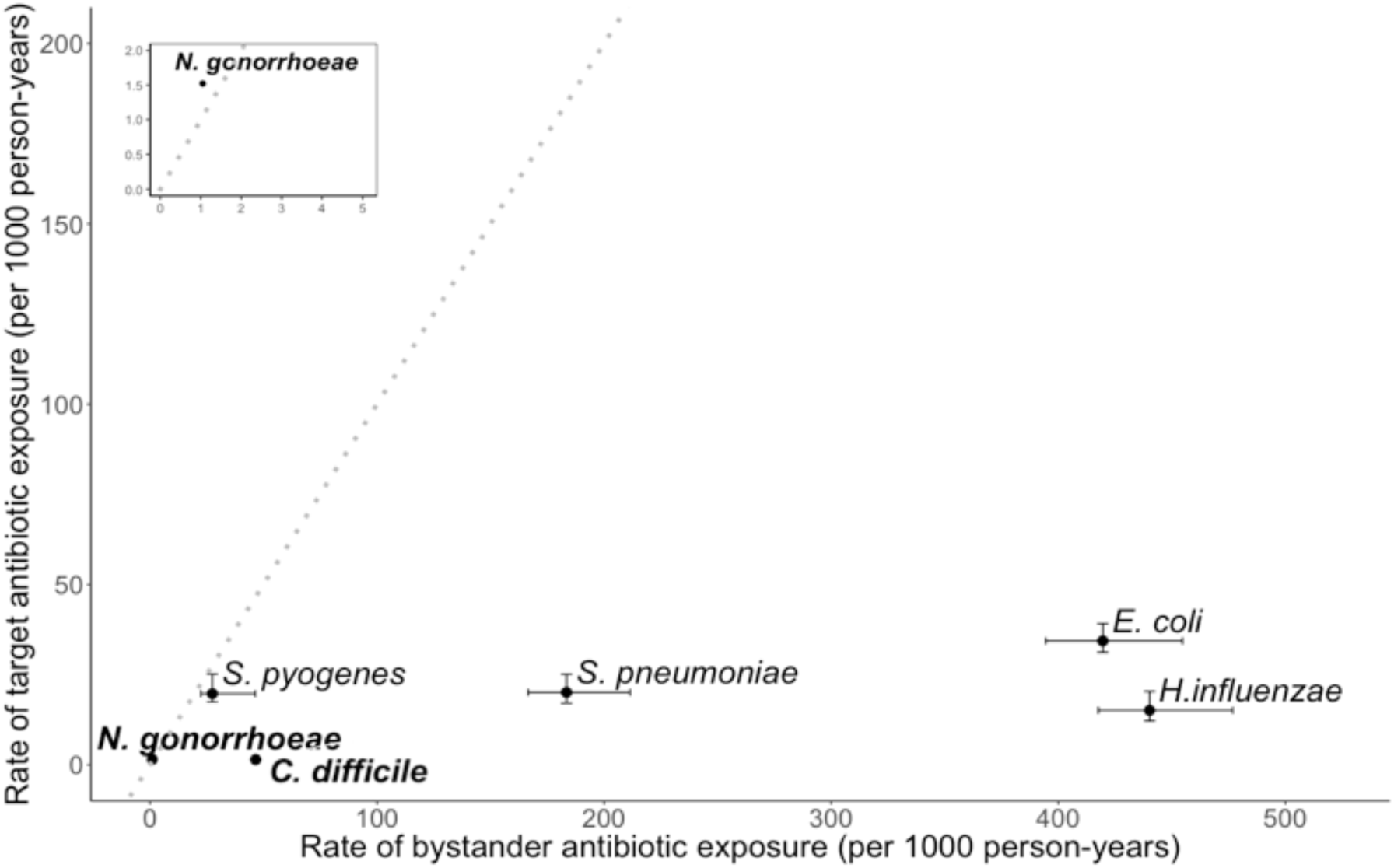
Incidental antibiotic exposure during asymptomatic carriage exceeds disease-related antibiotic exposure for key human pathogens. Bold font: Tier 1 urgent resistance concerns according to the Centers for Disease Control and Prevention (CDC) (1). Standard font: The most frequent etiologic agents of the top indications for antibiotic prescription in US ambulatory care. “Target antibiotic exposure” is defined as any antibiotic use associated with disease caused by that organism; “bystander antibiotic exposure” refers to the incidence of antibiotic exposure in asymptomatic carriage, roughly calculated as the product of the incidence of antibiotic prescription in ambulatory care and the proportion of the population in the relevant age group that carries the bacterium, less the number of target antibiotic exposures. The dotted line is where incidence of antibiotic exposure in carriage is equal to incidence of antibiotic exposure due to disease. See Tedijanto et al. (31) for method details, source references, and an alternate visualization of the same data on *N. gonorrhoeae, S. pyogenes, S. pneumoniae, E. coli*, and *H. influenzae*. Values for *C. difficile* were calculated using the same methodology and additional sources for disease incidence (33) and carriage prevalence (34); see SI.F for details.

An implication of our analysis is that, for pathogens like *S. pneumoniae, E. coli, H. influenzae*, and *C. difficile* that overwhelmingly face bystander exposure to antibiotics, even the strongest possible medical interventions informed by point-of-care resistance diagnostics can be insufficient to halt the rise of resistance to any drug in routine use. Mathematical details underpinning this analysis, including derivations of each strain’s reproduction number with and without POC-RD, are provided in SI.C.

In Fig 4, we parameterize our SCIS model for the pneumococcus, to show the threshold strain-1 fitness cost 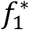 (contour lines) for a population with substantial bystander exposure (the *I*1 resistant strain is counter-selected in the blue regions above the contour lines). Pneumococcal serotypes show considerable variation in carriage duration (Fig 4, arrows on *x-*axis) (35) and, unsurprisingly, our analysis illustrates that bystander selection becomes increasingly problematic for longer carriage serotypes – represented by a higher threshold resistance cost 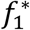 to balance selection on resistance. Our parameterized model indicates that the threshold costs even in the absence of RD are of a similar magnitude to *average* costs of plasmid encoded resistance discussed earlier (30). However, more problematic is the common observation of cost-free resistances (including single SNPs) in the pneumococcus (36, 37), indicating that these average costs are liable to decrease in response to selection.

**Fig 4.**
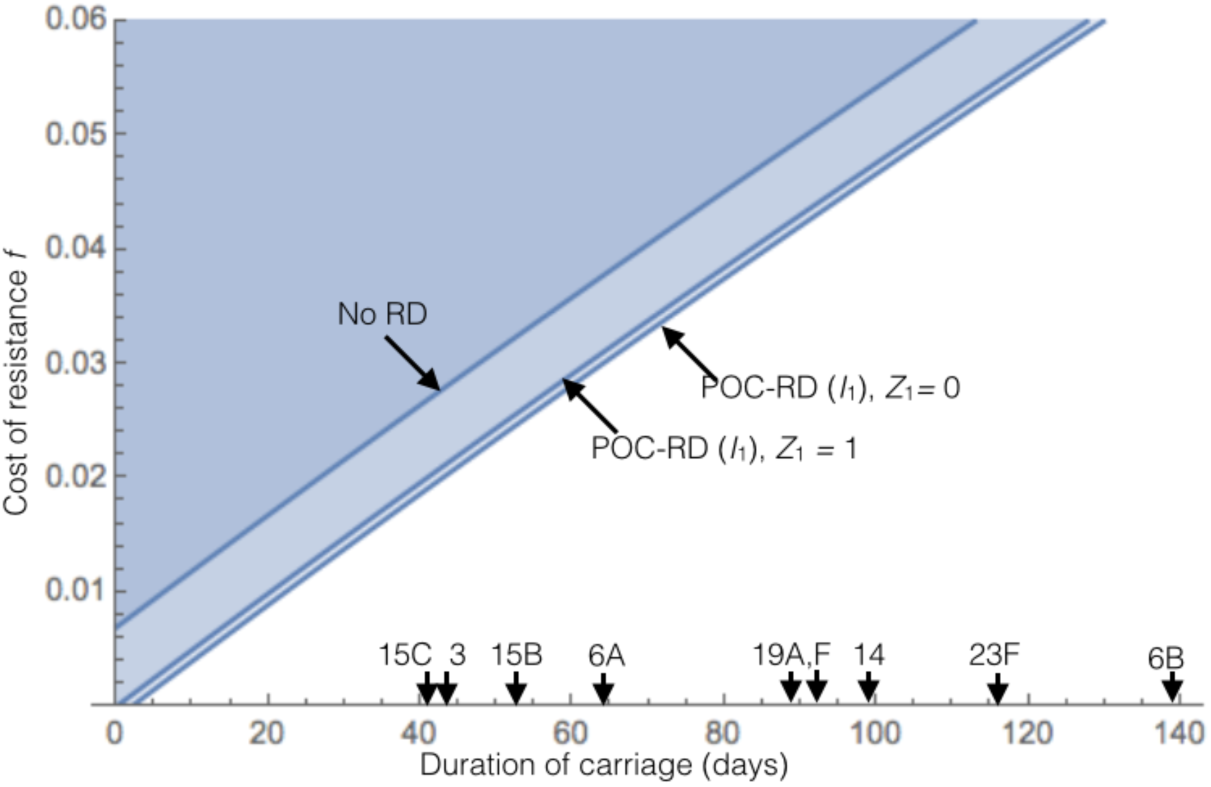
POC-RD alone cannot reverse selection on cost-free pneumococcal resistance. The minimal cost of resistance 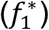 that allows universal treatment without causing an increase in strain-1 resistance is plotted (contour lines) against the expected duration of carriage. Two POC-RD scenarios are shown: with 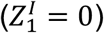 and without 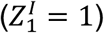 transmission control. Arrows on the *x* axis are serotype specific mean carriage duration estimates from (35) (serotypes with *≥15* recorded carriage episodes only). The remaining parameters (rates per day) are 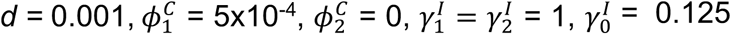. We make the simplifying assumption that baseline carriage and infection transmission rates are identical (*β*^*c*^ = *β*^1^ = *β*) given which 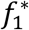 does not depend on *β*. Details on parameterization are in SI.G.

Even in the absence of pan-resistant strains, POC-RD has a weak impact due to the rarity and brevity of infection events (median infection duration in the absence of treatment is 8 days) relative to long periods of carriage and associated bystander selection. As a result, we anticipate ongoing selection for low-cost resistant strains, with or without POC-RD-informed strategies (Fig 4).

### Case #3: carriage resistance diagnostics

Our pessimistic conclusion concerning the public health merits of POC-RD for commensal opportunists such as *S. pneumoniae* (Fig 3,4) is based on the inevitability of bystander selection during prolonged carriage phases—but what if bystander selection could be opposed by public health interventions during carriage? For example, what if resistant-strain carriers could be identified and subjected to transmission control interventions even when they do not have active infection? Consider the South Swedish Pneumococcal Intervention Project (SSPIP) (38, 39), a public health intervention launched in January 1995 in Malmöhus County, Sweden that aimed to reduce penicillin-resistant pneumococcus (PRP) transmission, especially at preschool daycares. Any time a preschool-age child was identified with symptomatic PRP infection, providers would obtain nasopharyngeal cultures from all other children in the same daycare classroom. Children found to be carrying PRP were then required to remain home until subsequent testing proved them to be PRP-negative, penalizing PRP strains by reducing their opportunities for transmission from carriage.

In SI.C, we extend our analysis of the SCIS model to examine the potential of carriage RD-based interventions to generate selection against resistance. A key result is that, in order to select against drug-1-resistance, the drug-1-resistant strain must be discovered while in carriage at a rate *r*_1_ that exceeds the rate 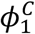 at which the sensitive strain is cleared from carriage due to bystander exposure to drug 1. In Fig 5, we again parameterize our SCIS model for the pneumococcus as in Fig 4, under two extreme scenarios of carriage duration (35): 20 weeks (Fig 5A) and 2 weeks (Fig 5B). The blue parameter space in Fig 5 highlights the combinations of carriage discovery rate (*r*_1_, x-axis) and HTC effectiveness against drug-1-resistant bacteria discovered in carriage (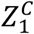 y-axis) that lead to a net selection against drug-1 resistance.

**Fig 5.**
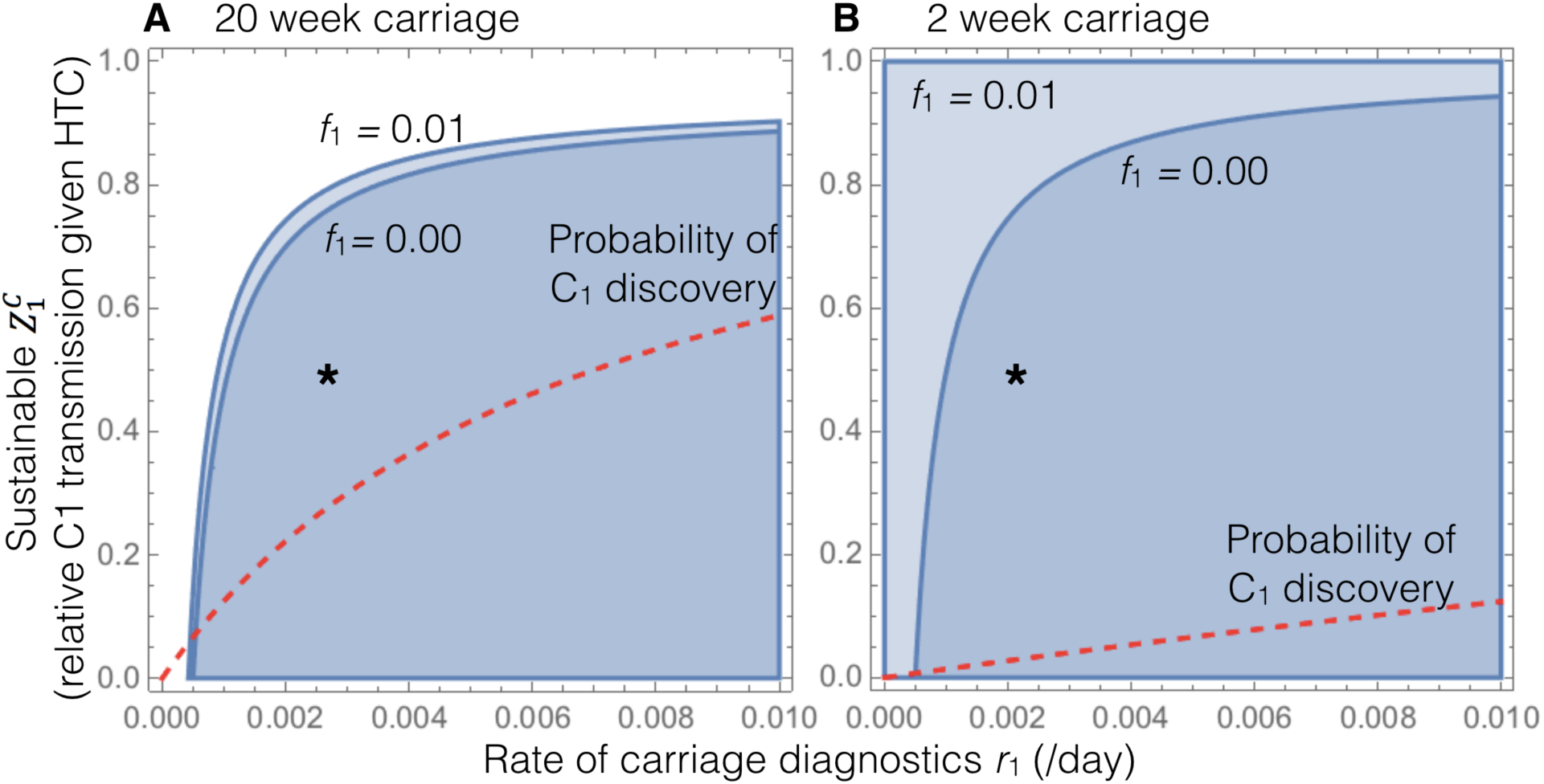
POC-RD plus Carriage RD can reverse selection on pneumococcal resistance, even for long carriage-duration serotypes. The parameter space generating net selection against resistance is plotted in blue as a function of the rate of carriage discovery (*r*_1_) and the effectiveness of carriage HTC 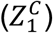. (A), longest carriage-duration serotype (6B, median 20 weeks). (B), shortest duration-carriage serotypes (1,4,5; ∼2 weeks). In both (A) and (B), two POC-RD scenarios are shown: with (*f* = 0.01) and without (*f* = 0) biological cost of resistance. The red dashed line represents the probability of strain 1 discovery while in the carriage state (“ *C*_1_ discovery”), an increasing function of the rate of carriage diagnosis. The remaining parameters (rates per day) are 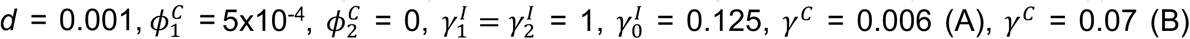. We make the simplifying assumption that baseline carriage and infection transmission rates are identical (*β*^1^ = *β*^0^ = *β*), given which the space of parameter generating net selection against resistance does not depend on *β*. Details on parameterization are in SI.G. The asterisk positions the outcome of an annual intervention with 50% efficacy in reducing *C*_1_ transmission.

Fig 5A illustrates the most problematic serotype from a POC-RD perspective, due to the dominance of bystander selection. Given the introduction of annual carriage surveillance (ensuring 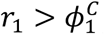), our parameterized model predicts that HTC interventions (such as removal of an infant from nursery) would need to reduce strain-1 transmission from carriage by at least 20% in order to select against the drug-1-resistant genotype in the worst case scenario of zero resistance costs (*f*_1_*=* 0). Epidemiological studies of the impact of child-care attendance on pneumococcal carriage suggest that a 2-fold reduction in transmission due to removal from daycare is not unreasonable (40).

The analysis underlying Fig 5 in SI.C implicitly assumes that all uninfected hosts are tested for drug-1-resistant carriage at a constant rate. While this assumption is useful here from an expositional point of view to highlight ideas, we note that such an approach would be inefficient in practice since many hosts would be tested even when their likelihood of drug-1-resistant carriage is low. By contrast, in SSPIP, only classmates of children who developed PRP infections were tested. Thus, even if PRP were rare in the general population, each child tested through SSPIP would have a substantial likelihood of PRP carriage. By introducing simple contact-tracing principles, the effective rate of carriage discovery can be higher for a given level of investment in patient sampling.

## Discussion

The antibiotic-resistance crisis is placing increasing pressure on healthcare globally and is widely viewed as a one-way street toward a dangerous “post-antibiotic world” (1, 4). In this paper, we ask whether resistance diagnostics (RD), when combined with public health interventions such as heightened transmission control (HTC) for drug-resistant bacterial strains, can substantially change the trajectory of resistance evolution.

In the early decades of the antibiotic era, doctors had no choice but to treat and control infection *unconditionally*, creating a selective pressure favoring strains that were resistant to whatever antibiotic was being widely prescribed. In that context, once resistant strains emerge with zero biological fitness costs, all antibiotics become “exhaustible resources” whose value to society is diminished by use (2, 3). In this paper, we show that exhaustible resource antibiotics can in principle be transformed into “renewable resources” whose value to society can be maintained over time even as they are put to widespread use, so long as (i) RD is available to detect the target pathogen and determine its antibiotic-sensitivity profile, (ii) prescribers adopt RD-informed treatment strategies, (iii) identified resistant cases are subject to more stringent transmission control, and (iv) bystander selection on the target pathogen is either minimal (Fig 2) or counteracted (Fig 5). For obligate pathogens that face minimal bystander selection, POC-RD-informed treatment reduces the advantage that resistant strains would otherwise enjoy (41) but, as we show, RD-informed treatment plus heightened control can potentially be sufficient to create a net selection against resistant pathogen strains, reducing their prevalence over time in the pathogen population. However, for opportunistic pathogens that face extensive bystander selection, POC-RD alone is insufficient—identifying resistant strains when they are not yet causing infection (so-called “carriage RD”) is essential to reverse the rise of resistance. This note of caution is important as society seeks to allocate resources most effectively in the struggle against antibiotic resistance.

Our analysis identifies strategies to renew or maintain sensitivity to an antibiotic in a pathogen population, the most effective of which depend on the availability of other antibiotics that can be used to treat resistant infections. The potential to restore antibiotic sensitivity is therefore limited once pan-resistant strains are in circulation. Consider now the impact of the discovery of a new antibiotic (drug 3) to which these bacteria are still sensitive. Drug 3’s discovery transforms previously pan-resistant bacteria into treatable “multidrug-resistant bacteria.” Providers can now deploy targeted treatment and HTC to hold the multidrug-resistant strain at a reproductive disadvantage. In this way, introducing a new antibiotic to which disease-causing strains are not yet resistant may make it possible to reverse the rise of previously pan-resistant bacteria, restoring the effectiveness of pre-existing antibiotics. Moreover, as multidrug resistance to pre-existing antibiotics grows less prevalent over time, the number of patients who need the new antibiotic will itself decline over time, allowing the new antibiotic to be held in reserve for increasingly rare cases for which it is the only effective treatment. This potential decline in drug 3 use highlights the importance of having drug 3 resistance diagnostics available when drug 3 is introduced, when we envisage the greatest use. Indeed, if drug 3’s introduction is coupled with drug 1-3 resistance diagnostics, it may not be necessary to discover even more new antibiotics *beyond* drug 3, as diagnostics will allow ‘search and destroy’ tactics against rare 1-3 pan resistant strains. We note that rolling out drug 3 at the same time as a drug 3 diagnostic raises the challenge of diagnosing resistance before widespread clinical use and clinical resistance discovery, necessitating increased investment in resistance discovery in the laboratory and phenotypic resistance surveillance in the clinic.

Whether net selection against resistant strains can be maintained depends on the effectiveness of the HTC measures that can be feasibly targeted against each resistant strain. By design, HTC measures impose additional barriers to resistant-bacterial transmission by (i) identifying resistant bacteria (during infection and/or asymptomatic colonization) and (ii) deploying additional resources specifically to prevent their transmission. Many sorts of HTC measures could be relevant in different contexts. Some examples: for hospital-associated infections, imposing heightened contact precautions when a hospitalized patient is found to have resistant infection (19, 21); for pneumococcal infection, requiring young children found to be infected or colonized with penicillin-resistant pneumococci to stay home from daycare (38); for sexually transmitted diseases, providing expedited partner therapy (EPT) when a patient is found to have resistant infection (42); or, for livestock-associated infections, eradicating an entire herd when resistant infection is identified. Note that HTC effectiveness may vary depending on the resistant strain being targeted, e.g., EPT measures may be less effective against pan-resistant strains since partners’ transmissibility cannot be controlled through treatment, while other measures may only be economically feasible against some strains, e.g., eradicating an entire herd may only be economical when pan-resistant infection is found, since other strains can be controlled through RD-informed treatment. More research is needed to quantify the effectiveness of HTC measures in practice.

Additional strategic options can be used to reduce the prevalence of bacterial strains with intermediate resistance, if RD is available that provides quantitative (e.g., sequence-based inference of minimal inhibitory concentration (43)) information on the degree of intermediate resistance. Details are provided in SI.B but, to see the point, imagine that POC-RD had been available when penicillin was first introduced that could quantitatively determine the penicillin sensitivity of gonorrhea infections. *Neisseria gonorrhoeae* strains emerged in the 1940s and 1950s that were less sensitive to penicillin but, at the time, these strains could still be effectively treated with a higher dose (44). Armed with quantitative POC-RD, providers would have been able to target intermediate-resistant gonococci with a higher penicillin dose—taking away the treatment-survival advantage that intermediate-resistant gonococci would otherwise enjoy—and could also have deployed additional public health resources to find and treat others who might still be spreading intermediate-resistant gonococci. Such a policy of *RD-informed treatment and heightened discovery* could have potentially held intermediate-resistant gonorrhea strains at an overall reproductive disadvantage relative to highly-sensitive strains, though only with HTC would it have been possible to avoid rapid selection of the highly resistant strains (45). See SI.B for mathematical details.

The example of the gonococcus raises the key challenge of bystander exposure to antibiotics, as gonorrhea infection is initially (and in some, entirely) asymptomatic and therefore does not drive immediate medical attention and exposure to POC-RD. During the asymptomatic phase of infection, drug-sensitive gonococcal genotypes are at risk of being cleared due to antibiotics taken for other medical concerns (46). In Fig 3, we outline how the extent of the bystander challenge is even greater for commensal opportunistic pathogens (47, 48) which spend proportionately longer in asymptomatic carriage states. Parameterizing our SCIS model (incorporating a carriage / asymptomatic stage, prone to bystander selection) for the key commensal opportunist *S. pneumoniae* illustrates that selecting against resistance via POC-RD-informed strategies alone is not a plausible outcome for this particular pathogen (Fig 4), given the lengthy duration of carriage and relatively rare and brief infection events caused by this species.

We explore a strategic response to this concern: conditional interventions in response to diagnostic information during asymptomatic carriage. Fig 5 illustrates that coupling differential transmission control to carriage RD can drive net selection against resistance, even for the most carriage-prone serotypes of the pneumococcus. The South Swedish Pneumococcal Intervention Project (SSPIP) offers a concrete example of using carriage RD to drive public health interventions. While SSPIP targeted pneumococcal strains that remained treatable by other antibiotics, similar programs could target pan-resistant strains and, if sufficiently intensive and comprehensive, potentially select against these pan-resistant strains even as those with sensitive infection continue to receive antibiotic treatment.

We note that, in theory, our logic of conditional interventions during carriage could be extended to incorporate broader ‘microbiome’ resistance diagnostics (M-RD) and M-RD-informed interventions. While simple in outline, implementation presents technical challenges on several fronts, not least in establishing meaningful sampling protocols, designing appropriate narrow-spectrum interventions (10, 49–54), and designing appropriate strategic rules for intervention choice given potentially conflicting microbiome and infection-site resistance profiles. We also note that, independent of any M-RD innovations, the widespread uptake of POC-RD-informed antibiotic use will likely reduce bystander selection due to an overall reduction in antibiotic use (e.g., in the context of viral infections) and a potential shift toward narrower-spectrum antibiotics being prescribed against known pathogen targets.

Our focus on homogenous, closed populations ensures that R_0_ maximization is always favored by selection, simplifying the evolutionary analysis (55). However, the basic idea underlying our analysis of modulating strain-fitness landscapes applies more broadly to models where R_0_ is not a sufficient proxy for fitness, such as in cases of multiple carriage (55), host population structure (56), or an open population (28).

In the supplementary material, we extend our SIS model analysis to consider the effect of antibiotic rationing whereby some patients are left untreated (SI.B) and show that our findings are robust to (i) environmental reservoirs of resistant bacteria (SI.D), (ii) host migration from high-resistance regions (SI.D), (iii) resistance-conferring mutation (SI.D), (iv) competitive release (SI.D), (v) diagnostic errors (SI.E), and (vi) diagnostic escape (SI.E). Inflows of resistant cases (mutation, migration) together with diagnostic errors weigh on the scale in favor of resistant strains but can all be counteracted by sufficiently high reproductive penalties to correctly-targeted resistant strains. Diagnostic escape (57) presents a qualitatively distinct challenge where diagnostic tests themselves become obsolete due to evolutionary responses in the pathogen (e.g., loss or modification of resistance marker). The risk of diagnostic escape highlights the importance of active resistance surveillance and rapid new-diagnostic development.

Point-of-care resistance diagnostics are already a top public-health priority, with a major emphasis on rapidity (<1 hour) (58). Rapid point-of-care diagnostics are critical for early effective treatment of life-threatening infections when treatment cannot be delayed. However, most antibiotic prescriptions are for less severe infections where patients can wait longer to benefit from more complete diagnostic information (13). Our analyses illustrate that, for the public health goal of selecting against resistance in pathogens with minimal bystander selection, we have more time to act—delays until treatment on the order of hours or even days following initial infection may still allow for selection against resistance (Fig 2), However, our conclusions depend critically on the life-history of the target pathogen, with the message that reversing resistance in opportunistic pathogens subject to ‘bystander’ selection is not generally plausible with POC-RD information alone (Fig. 4), and will require additional interventions conditioned on carriage RD (Fig 5). Our conclusions also depend on the ability of RD to distinguish multiple resistances in a multi-drug context, highlighting the importance of diagnostic breadth as well as rapidity. Diagnostic-informed approaches to reversing resistance face another time constraint—our proposed strategies for resistance-targeted intervention are most effective when pan-resistant strains are still rare (SI.B). If we fail to act decisively while bacteria that are resistant to all antibiotics remain rare (59, 60), we may then be unable to reverse the continued rise of untreatable bacterial disease.

## Supplementary Information

is available in the online version of the paper.

### Acknowledgements

We thank Luke McNally, Dan Cornforth, Steve Diggle, Arjun Srinivasan, Cliff McDonald and Alison Halpin for discussion and comments on earlier drafts. The project described was supported by the CDC (OADS BAA 2016-N-17812), the National Institute of General Medical Sciences (U54GM088558), the Simons Foundation (396001), HFSP (RGP0011/2014), the Wenner-Gren Foundations, and the Royal Physiographic Society of Lund. The content is solely the responsibility of the authors and does not necessarily represent the official views of the CDC, the National Institute of General Medical Sciences or the National Institutes of Health, or any other supporting organization.

## Author Contributions

DM conceived the project. DM and SB developed the mathematical models. DM analyzed the models. KWW, CT & ML reviewed and summarized epidemiological data. DM and SB wrote the manuscript. DM wrote SI.A-E, CT wrote SI.F, and KWW wrote SI.G. All authors edited the manuscript and discussed the modelling approach.

