## Supplementary Information for "Resistance diagnostics as a public health tool to combat antibiotic resistance: A model-based evaluation"

This supplement provides supporting discussion and mathematical detail.

Part A is for readers interested in more non-mathematical discussion of the paper’s main thesis that, in some cases, resistant-pathogen strains can be held at a selective disadvantage through resistance diagnostic (RD)-informed interventions. To keep the analysis as clear and simple as possible, the manuscript (MS) focuses on a specific context in which providers and public-health authorities’ only non-antibiotic intervention option is to deploy “heightened transmission control (HTC)” measures that reduce the transmissibility of identified resistant-pathogen carriers. Part A provides further discussion of HTC-oriented strategies but also discusses other sorts of possibilities, such as RD-informed contact tracing, that can accelerate resistant-pathogen discovery.

Parts B-C are for readers interested in mathematical details supporting the analysis in the MS and Part A. Part B provides detailed derivations of the mathematical expressions in MS Case #1 on an obligate pathogen while Part C provides details for Cases #2-3 on an opportunistic pathogen. In addition, Part B extends the obligate-pathogen analysis in two new directions discussed in Part A, allowing for (i) RD-informed discovery efforts referred to as “heightened discovery (HD)” and (ii) rationing of antibiotic treatment, relaxing the assumption in the MS that all patients must receive the most effective treatment.

Parts D-E explore the robustness of our main findings in a variety of extensions. Part D extends the analysis to allow for inflows of resistant pathogens due to resistance-conferring mutation, migration of infected hosts, and environmental exposure to resistant pathogens, while Part E allows for diagnostic errors and diagnostic escape.

Part F provides details on data sources for Fig 3, which is based on a similar figure in Tedijanto et al (2018) but (i) provides a different visualization of the data that is more useful for motivating this paper’s analysis and (ii) includes data for *C. difficile*.

Part G provides details supporting the parameter values used to construct Figs 4-5.

### Part A. Verbal/schematic summary: Resistance-targeted intervention

This section provides an expanded informal discussion of the basic idea underlying our analysis, that public-health interventions aimed at more-effectively discovering, treating, and controlling resistant strains can reduce and in some cases reverse resistance-favoring selection.

1. Resistance-targeted antibiotic treatment

In a world where all infections are treated with a first-line antibiotic (drug 1), bacteria that acquire drug-1 resistance will enjoy a powerful reproductive advantage, whether or not they are resistant to other antibiotics. Once point-of-care resistance diagnosis (POC-RD) is available, however, infections that are resistant to drug 1 but sensitive to some other antibiotic ("drug 2") can be effectively treated with that other antibiotic. Simply acquiring drug-1 resistance is therefore no longer enough for bacteria to enjoy a reproductive advantage. Indeed, so long as (i) drug 2 is equally effective against drug-1-resistant bacteria as drug 1 is against drug-1-sensitive bacteria and (ii) there are biological fitness costs associated with drug-1 resistance, bacteria that are resistant to drug 1 but not drug 2 will be at a strict reproductive disadvantage relative to those that are sensitive to both drugs.

What if drug 2 is less effective than drug 1 (or drug 1 is the only treatment option)? Drug-1-resistant bacteria treated with drug 2 (or left untreated) will then enjoy a reproductive advantage over drug-1-sensitive bacteria treated with drug 1 (Fig. SI.1b-c), and we can expect drug-1-resistant infection to grow more prevalent over time even when POC-RD is available. Fortunately, there are several ways potentially to counteract this remaining advantage, such as identifying resistant infections more quickly ("resistance-targeted discovery") and intensifying efforts to reduce resistant-bacterial transmission opportunities ("resistance-targeted infection control" in the hospital context and/or "resistance-targeted social distancing" in the community context).

2. Resistance-targeted discovery

When a patient is found to have drug-1-resistant infection, hospital administrators (for hospital-associated infections) and public-health authorities (for community-associated infections) have more incentive than otherwise to make an effort to identify others with drug-1-resistant infection. For instance, in the case of gonorrhea, suppose that POC-RD were available to detect ciprofloxacin resistance. Patients found to have ciprofloxacin-sensitive infection could be treated with ciprofloxacin, while those found to have ciprofloxacin-resistant infection could be treated with the alternative treatment of azithromycin and ceftriaxone (1). Armed with the knowledge of who has ciprofloxacin-resistant infection, public-health officials could then work even more intensively to identify at-risk sexual partners, to discover others infected with the ciprofloxacin-resistant strain who have not yet sought out treatment, and accelerate their recovery by treating them with the azithromycin-ceftriaxone combination. So long as ciprofloxacin-resistant gonorrhea remain sensitive to the azithromycin-ceftriaxone combination, such accelerated treatment can put ciprofloxacin-resistant strains at a reproductive disadvantage relative to ciprofloxacin-sensitive strains that do not trigger heightened discovery (Fig. SI.1c-d).

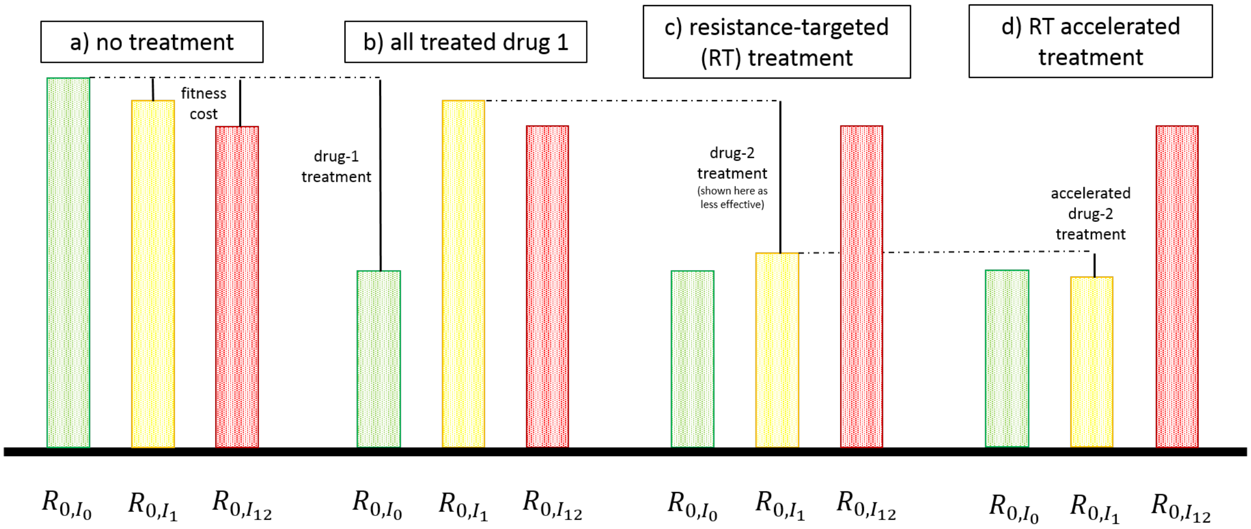


**Figure SI.1**. **Resistance-targeted treatment.** Reproduction numbers of sensitive strain ($I_0$), drug-1-resistant strain ($I_1$), and pan-resistant strain ($I_{12}$), under various medical interventions: **a)** *no treatment*, $I_0$ enjoys advantage due to fitness costs associated with resistance; **b)** *all treated with drug 1*, $I_1$ and $I_{12}$ enjoy advantage due to drug-1 resistance; **c)** *resistance-targeted treatment*, $I_1$ now at disadvantage unless (as shown) drug 2 is sufficiently less effective than drug 1; and **d)** *resistance-targeted accelerated treatment*, whereby strain-$I_1$ infections come more quickly to medical attention due to heightened discovery.

What if pan-resistant bacteria have already emerged that cannot be treated by any antibiotic? Pan-resistant strains can still potentially be held at a reproductive disadvantage through infection-control efforts that more intensively disrupt pan-resistant transmission during infection or, at least in principle, by intervening prior to the onset of infection through “carriage treatments” that reduce transmissibility and/or clear pan-resistant bacteria while they remain in the asymptomatic carriage state.

3. Resistance-targeted infection control and/or resistance-targeted social distancing

Suppose for a moment that infected patients can be isolated sufficiently effectively to put isolated pan-resistant bacteria at a reproductive disadvantage relative to unisolated sensitive bacteria treated with drug 1. Once armed with POC-RD, doctors could then put pan-resistant bacteria at a reproductive disadvantage relative to sensitive bacteria—even while prescribing drug 1 to all those who would benefit from it—by isolating those with pan-resistant infection (Fig. SI.2a-b). Intuitively, targeted isolation creates a new “fitness cost” associated with pan-resistance, since only pan-resistant bacteria face isolation.

When isolation or other sufficiently effective infection-control options are available, pan-resistance-targeted infection control can be enough to reverse the rise of pan-resistance. On the other hand, when infection-control options are less effective, the “fitness cost” that they impose is not enough, on its own, to put pan-resistant bacteria at a reproductive disadvantage (Fig. SI.2c). In such cases, accelerated discovery of pan-resistant infections could in principle amplify the impact of pan-resistance-targeted infection control (Fig. SI.2d). For example, consider a hypothetical bacterial disease that is sufficiently mild that most infected hosts never seek medical attention but, due to concerns about rising pan-resistance, public-health authorities conduct contact tracing from any index patient found to pan-resistant infection and impose heightened transmission control (HTC) measures (such as requiring children to stay home from school) on anyone found to have pan-resistant infection. If each such contact-tracing effort identifies on average two pan-resistant cases that never otherwise would have come to medical attention, the impact these extra control measures have on the pan-resistant strain’s reproduction number will be three times larger than if resistance-targeted contact tracing had not been conducted.

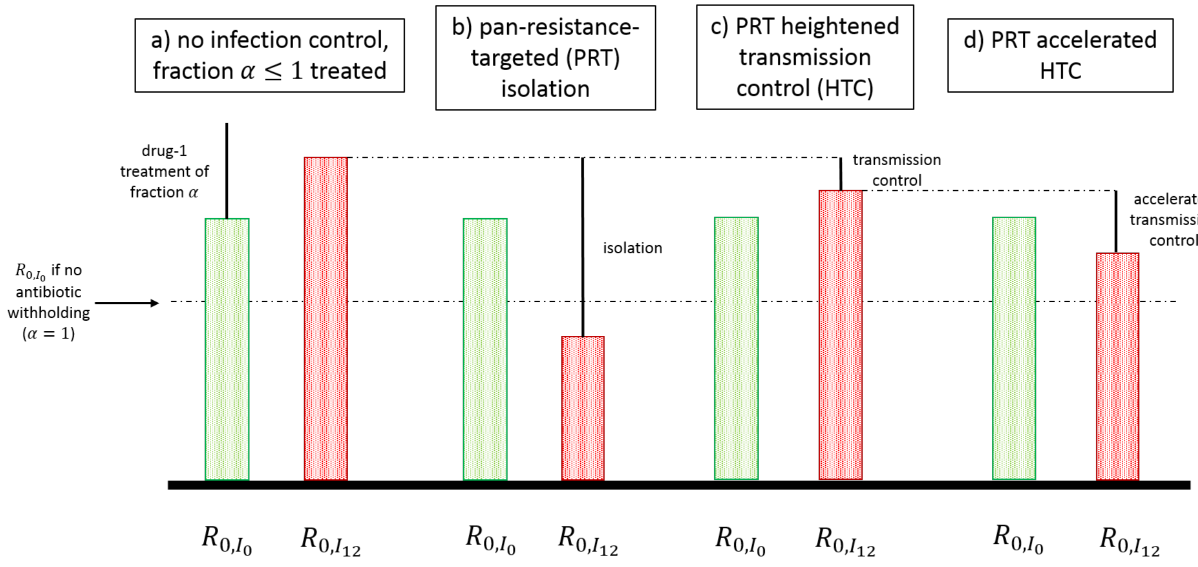


**Figure SI.2**. **Resistance-targeted infection control.** Reproduction numbers of sensitive strain ($I_0$) and pan-resistant strain ($I_{12}$), under various medical interventions: **a)** *fraction* $\alpha \leq 1$ *treated with drug 1*, $I_{12}$ enjoys advantage due to surviving treatment unless $\alpha$ is sufficiently small; **b)** *pan-resistance-targeted (PRT) isolation*, $I_{12}$ now at disadvantage even if all sensitive infections treated with drug 1; **c)** *PRT heightened transmission control (HTC)*, with HTC shown here as much less effective than isolation; **d)** *PRT accelerated HTC*, whereby strain-$I_{12}$ infections come more quickly under control due to heightened discovery.

4. Resistance-targeted carriage intervention

Many of the most important disease-causing bacteria are opportunistic pathogens that dwell for extended periods in an asymptomatic carriage state. Bystander antibiotic exposure in carriage gives resistant strains a reproductive advantage, even if antibiotic treatment is withheld during infection (Fig. SI.3a). Moreover, for pathogens that dwell mainly in carriage, the effect of bystander antibiotic exposure may be too large to be overcome by any point-of-care intervention on the treatment of infections they cause (Fig. SI.3b; see Part C for details). In such cases, the only way to put resistant bacteria at a reproductive disadvantage is to intervene in carriage, identifying when patients are colonized with a resistant strain—*before* the emergence of harmful symptoms—and then treating and/or controlling resistant bacteria found in carriage (Fig. SI.3c).

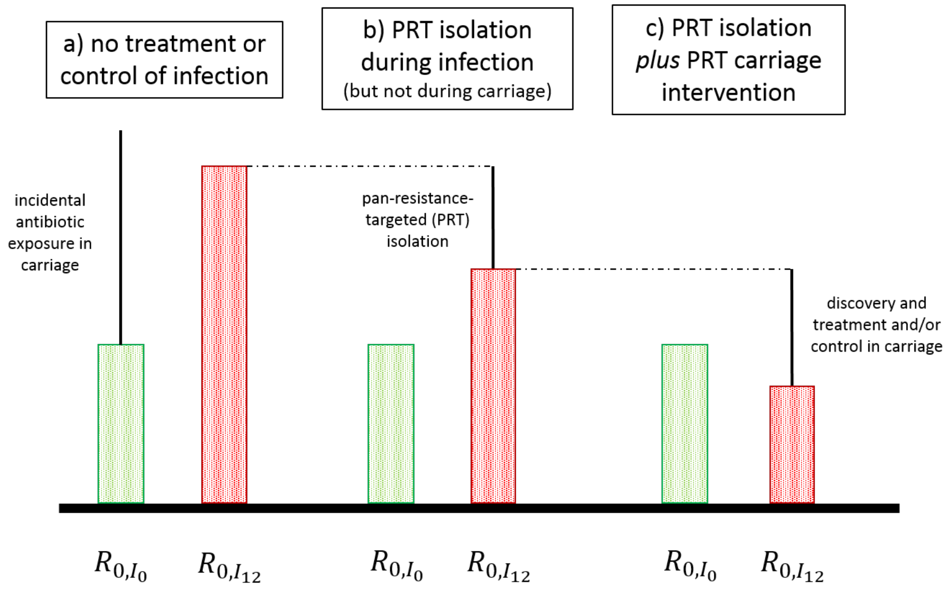


**Figure SI.3**. **Resistance-targeted carriage intervention.** Reproduction numbers of sensitive strain ($I_0$) and pan-resistant strain ($I_{12}$) of an opportunistic pathogen, under various medical interventions: **a)** no treatment or control, $I_{12}$ enjoys advantage due to surviving incidental antibiotic exposure; **b)** pan-resistance-targeted (PRT) isolation during infection, $I_{12}$ may still enjoy advantage if the pathogen dwells mainly in carriage; **c)** PRT isolation during infection plus PRT carriage intervention, $I_{12}$ now at overall disadvantage so long as carriage intervention more effective at clearing pan-resistant bacteria than incidental exposure is at clearing sensitive bacteria (see Part C for details).

*Discussion: carriage treatments*. Pan-resistant bacteria cannot be effectively controlled while causing infection, but in some cases there may be "carriage treatments" to which they are still susceptible. For instance, suppose that an *E. coli* strain has been engineered that can sense and kill a pan-resistant strain of the pathogen by producing pathogen-specific bacteriocins (2, 3) If such killer-*E. coli* treatment is not fast-acting enough to provide relief during infection, it could still be prescribed against pan-resistant bacteria discovered in carriage. So long as such treatment is prescribed sufficiently often against the pan-resistant strain while in carriage—and not prescribed (or prescribed sufficiently less often) when sensitive strains of the pathogen are discovered in carriage—the pan-resistant strain can be held at an overall reproductive disadvantage.

*Discussion: transmission control during asymptomatic carriage*. Carriage treatments are not yet widely available, but traditional infection-control methods can achieve the same effect if resistant-strain carriers can be identified and socially isolated even when they do not have active infection. In the manuscript, we discuss one such effort, the South Swedish Pneumococcal Intervention Project (SSPIP)

(4, 5), a public-health intervention launched in January 1995 in Malmohus County, Sweden that reduced transmission of penicillin-resistant pneumococci (PRP) at preschool daycares by requiring PRP carriers to remain home until proven to be PRP-negative.


### Part B. An obligate pathogen

Here we consider the case of an obligate pathogen, assuming for simplicity that only two antibiotic-treatment options are available (first-line therapy “drug 1” and second-line therapy “drug 2”), resistance to the first-line therapy has emerged, pan-resistance may or may not have emerged, and POC-RD is available to immediately determine the bacterial strain causing each patient’s infection.

This section is organized as follows. We begin by analyzing an especially simple “benchmark model” in which the formula for each strain’s reproduction number is easy to derive. We then extend that basic analysis in several ways, allowing for the possibility that (i) pathogen strains may exhibit intermediate resistance (mentioned in the Discussion section of the MS), (ii) there may be diagnostic delay (the main case analyzed in the MS), (iii) infected hosts may not be immediately brought to medical attention and public-health efforts (termed “heightened discovery (HD)”) may allow resistant infections to be identified more quickly, and/or (iv) antibiotic treatment may be rationed.

Later sections provide additional extensions, allowing for asymptomatic colonization by an opportunistic pathogen (Part C), inflows of resistance due to mutation, migration, or environmental infection (Part D), and diagnostic failures due to diagnostic error or diagnostic escape (Part E).

**Benchmark model: obligate pathogen**

Bacterial transmission and host recovery are governed by a standard Susceptible-Infected (SIS) model with multiple competing bacterial strains, augmented by a model of POC-RD-targeted medical intervention.

*Host states*. There is a unit-mass population of hosts, each of whom transitions over time between the following host states: $S$ when uninfected and susceptible to infection; $I_0$ when infected with a strain sensitive to both drugs; and $I_1$, $I_2$, $I_{12}$ when infected with a strain resistant only to drug 1, resistant only to drug 2, or resistant to both drugs ("pan-resistant"), respectively. For each $X \in \{0,1,2,12\}$, the notation $I_X$ will also be used to denote the mass of hosts infected with "strain $X$," the bacterial strain resistant to all antibiotics in $X$ and sensitive to all antibiotics not in $X$. In the special case when pan-resistance has not yet emerged, the only bacterial strains in circulation are strains 0, 1, and 2.

*Transmission*. Each host with strain-$X$ infection transmits strain-$X$ bacteria to a new host at baseline transmission rate $\beta_X > 0$. Such transmission events lead to new infection if the exposed host is currently uninfected but otherwise have no effect. (For simplicity, we assume that the host cannot be colonized by multiple strains at the same time, i.e., there is no co-infection.) There may or may not be biological transmission-related fitness costs associated with resistance; in particular, we assume that $\beta_X = (1 - f_X)\beta$ for each resistant strain X = 0, 1, 2, 12, where $f_X \geq 0$ captures any biological fitness costs associated with resistance to drugs X and $f_0 = 0$. For notational simplicity, we drop all "I" superscripts here when analyzing the obligate-pathogen model.

*Discovery and diagnosis*. Each infected host immediately comes under medical attention. At the point of care, the care provider ("doctor") conducts point-of-care resistance diagnosis (POC-RD), decides what if any antibiotic treatment to prescribe, and decides what if any additional transmission-control measures to order. When making these decisions, the doctor is assumed to comply with whatever treatment-and-control policy has been designated as the standard of care.

*Treatment*. There are three options for antibiotic treatment: drug 1, drug 2, or none. Infections that are untreated or ineffectively-treated clear at rate $\gamma_0 > 0$, those effectively treated with drug 1 or drug 2 recover at rate $\gamma_1 > \gamma_0$ or $\gamma_2 \geq \gamma_0$, respectively. We focus on the case in which drug 1 is at least as effective as drug 2, i.e., $\gamma_1 \geq \gamma_2$.[1] (The special case in which $\gamma_2 = \gamma_0$ is equivalent to drug 1 being the only treatment option.) We focus on treatment policies in which all patients receive the most-effective antibiotic treatment, i.e., (i) those found to have strain-0 or strain-2 infection are treated with drug 1, (ii) those found to have strain-1 infection are treated with drug 2, and (iii) those found to have strain-12 infection are left untreated.

*Transmission control*. There are three options for transmission control: standard measures (or "no control"), heightened transmission control measures that can be targeted against resistant strain $X$ ($HTC_X$), or isolation. Isolation proportionally reduces transmission to a fraction $Z_{isolate} < \frac{\gamma_0}{\gamma_1}$ of what it would otherwise be; however, there is limited capacity to isolate at most mass $\epsilon \geq 0$ of infected hosts at any given time, out of an overall host population of mass one. (In the manuscript, we focus on the case in which patient isolation is unavailable, i.e., $\epsilon = 0$.) $HTC_X$ reduces transmission of strain-$I_X$ bacteria by factor of $Z_X < 1$ and can be ordered for an unlimited number of patients. The condition $Z_{isolate} < \frac{\gamma_0}{\gamma_1}$ implies that a host who is isolated and left untreated will generate fewer transmission events over the course of their infection than one who is treated with drug 1 and subjected to standard transmission control. $\epsilon = 0$ corresponds to a scenario in which infected hosts cannot be isolated, while $Z = 1$ corresponds to a scenario in which heightened transmission control is not possible. We focus on control policies in which (i) all strain-0 infections are subjected to standard measures, (ii) all strain-1 and strain-2 infections are subjected to HTC but not isolated, and (iii) all strain-12 infections are isolated as much as possible (until isolation capacity $\epsilon$ is filled) and otherwise subjected to HTC.

*Note on notation:* Depending on the context, subscripts indicate either the resistance profile of a bacterial strain (e.g., $\beta_1$ is the transmission rate of the drug-1-resistant strain 1) or the recovery rate associated with an antibiotic treatment (e.g., $\gamma_1$ is the recovery rate of any drug-1-sensitive infection treated with drug 1, while $\gamma_0$ is the recovery rate of any untreated infection).

*Discussion: heightened transmission control ($HTC_X$).* Resistance diagnosis makes it possible to reduce the transmissibility of resistant infections in many ways. Some examples: *Patient self-isolation*: Once

[1] All of our analysis extends in a straightforward way to the case in which drug 2 is more effective. Most notably, if $\gamma_2 > \gamma_1$, then the drug-1-resistant strain 1 can be held at a reproductive disadvantage relative to the sensitive strain 0 merely by treating drug-1-resistant infections with drug 2.

informed that they have resistant infection, patients can reduce their own transmissibility by washing their hands more frequently, not sharing towels, etc. *Socially-imposed isolation*: Patients found to be harboring resistant bacteria could be required to stay home from school or work until they can be confirmed non-contagious. *Vaccination of at-risk contacts*: Currently-uninfected family members and others close to a patient found to have resistant infection could receive vaccination or prophylactic treatment (if available) to reduce the frequency of bacterial transmission.

**Reproduction number of each strain**

Let $R_{0,X}$ denote strain $X$'s basic reproduction number (or simply "reproduction number"), the average number of transmission events during the course of strain-$X$ infection. ($R_{0,X}$ is the average number of additional infections caused by a single strain-$X$ infection in an otherwise completely-susceptible population.) A basic result in SIS models with competition for hosts is that whichever strain has the highest reproduction number will, in the long run, come to dominate the bacterial population, driving the burden of infection caused by the other strains to zero. We are interested in identifying conditions in which the sensitive strain enjoys a strict reproductive advantage over all resistant strains currently in circulation. Under such conditions, all resistant strains will grow less prevalent over time, i.e., "resistance is reversed."

_Sensitive strain:_ Strain-0 infections have average duration $\frac{1}{\gamma_1}$ because they are treated with drug 1 and transmit at rate $\beta_0 = \beta$ because they are subjected to standard control; so, $R_{0,0} = \frac{\beta}{\gamma_1}$.

_Drug-1-resistant strain:_ Strain-1 infections have average duration $\frac{1}{\gamma_2}$ because they are treated with drug 2 and transmit at rate $Z_1\beta_1 = (1 - f_1)Z_1\beta$ because they are subjected to HTC$_1$; so, $R_{0,1} = (1 - f_1)Z_1\frac{\beta}{\gamma_2}$.

_Drug-2-resistant strain:_ Strain-2 infections have average duration $\frac{1}{\gamma_1}$ because they are treated with drug 1 and transmit at rate $Z_2\beta_2 = (1 - f_2)Z_2\beta$ because they are subjected to HTC$_2$; so, $R_{0,2} = (1 - f_2)Z_2\frac{\beta}{\gamma_1}$.

_Pan-resistant strain:_ Strain-12 infections have average duration $\frac{1}{\gamma_0}$ because they are left untreated.

If pan-resistance is sufficiently rare that $I_{12} \leq \epsilon$, all strain-12 infections can be isolated. Otherwise, if $I_{12} > \epsilon$, fraction $\frac{\epsilon}{I_{12}}$ of strain-12 infections can be isolated while the rest can be subjected to HTC$_{12}$. Let

$\hat{Z} = \frac{\epsilon}{I_{12}} Z_{isolate} + \left(1 - \frac{\epsilon}{I_{12}}\right) Z_{12}$ if $I_{12} > \epsilon$ or $\hat{Z} = Z_{isolate}$ if $I_{12} \leq \epsilon$. By definition of $\hat{Z}$, $\hat{Z}\beta_{12}$ is the average transmission rate of strain-12 infection; so, strain 12's reproduction number is then $R_{0,12} = (1 - f_{12})\hat{Z}\frac{\beta}{\gamma_0}$. Let $I_{12}^* > \epsilon$ be the threshold for pan-resistant infection prevalence given which $\hat{Z} = \frac{\gamma_0}{\gamma_1}$, defined as

$$I_{12}^* = \frac{\epsilon(Z_{12} - Z_{isolate})}{Z_{12} - \frac{\gamma_0}{\gamma_1}}$$

*Equation 1*

So long as pan-resistant infection is sufficiently rare that $I_{12} < I_{12}^*$, isolating mass $\epsilon$ of pan-resistant-infected hosts and subjecting the rest to $HTC_{12}$ is sufficient to hold the pan-resistant strain at a reproductive disadvantage relative to the sensitive strain, even as all sensitive infections are treated with drug 1.

**Threshold fitness costs**

Let ${f_X}^*$ denote the "threshold fitness cost" for strain X, the smallest biological fitness cost given which strain X is not at a reproductive advantage relative to strain 0.

*Drug-1 resistance:* The drug-1-resistant strain is at a reproductive disadvantage relative to the sensitive strain whenever $R_{0,0} = \frac{\beta}{\gamma_1} > (1 - f_1)Z_1\frac{\beta}{\gamma_2} = R_{0,1}$, i.e., whenever $f_1 > f_1^* = 1 - \frac{\gamma_2}{Z_1\gamma_1}$. Note that the threshold fitness cost $f_1^* < 0$ so long as drug 2 is equally effective against drug-1-resistant infections as drug 1 is against sensitive infections ($\gamma_2 = \gamma_1$) and $HTC_1$ is even modestly effective ($Z_1 < 1$). Thus, under those conditions, selection against drug-1 resistance can be maintained even if there are no biological fitness benefits associated with drug-1 resistance.

*Drug-2 resistance:* The drug-2-resistant strain is at a reproductive disadvantage relative to the sensitive strain whenever $\frac{\beta}{\gamma_1} > (1 - f_2)Z_2\frac{\beta}{\gamma_1}$, i.e., whenever $f_2 > f_2^* = 1 - \frac{1}{Z_1}$. Note that the threshold fitness cost $f_2^* < 0$ so long as $HTC_2$ is even modestly effective ($Z_2 < 1$). So, selection against drug-2 resistance can be maintained even if there are no biological fitness benefits associated with drug-2 resistance.

*Pan resistance:* The pan-resistant strain is at a reproductive disadvantage relative to the sensitive strain whenever $\frac{\beta}{\gamma_1} > (1 - f_{12})\hat{Z}\frac{\beta}{\gamma_0}$, i.e., whenever $f_{12} > f_{12}^* = 1 - \frac{\gamma_0}{\hat{Z}\gamma_1}$. If pan resistance is sufficiently rare that it can be effectively isolated ($I_{12} < I_{12}^*$), $f_{12}^* < 0$ and the pan-resistant strain can be held at a reproductive disadvantage even if there are no biological fitness costs associated with pan resistance. On the other hand, should pan-resistant infection become sufficiently widespread that only a small fraction of pan-resistant infections can be isolated ($I_{12} \gg I_{12}^*$), their average transmission rate $\hat{Z} \approx Z_{12}$ and so the threshold fitness cost $f_{12}^* \approx 1 - \frac{\gamma_0}{Z_{12}\gamma_1}$.

### Extension: Intermediate resistance

In this section, we extend the analysis to allow for the possibility that bacterial strains may be somewhat but not completely resistant to antibiotic treatment ("intermediate resistance"). For simplicity, and to focus on the most challenging case, suppose that drug 1 is the only effective treatment, there are no fitness costs associated with drug-1 resistance, and bacterial strains are in circulation that are resistant to drug 1 to various degrees. Bacterial strains with intermediate resistance can in some cases be held at a reproductive disadvantage without needing to target such infections for HTC, if instead such strains can be targeted for heightened dosing and/or heightened patient compliance.

#### Intermediate resistance-targeted heightened dosing

Suppose for simplicity that each strain is characterized by the minimal inhibitory concentration (MIC) of drug 1 needed in inhibit in vitro growth and that a strain with MIC $M$ will clear at rate $\gamma(X; M)$ when treated with drug 1 at dose $X$, where the recovery-rate function $\gamma(X; M)$ is continuous in $(X, M)$, increasing in $X$, and decreasing in $M$ with $\gamma(0; \infty) = \gamma_0$ the untreated recovery rate. Let "strain 0" be the most-sensitive strain (or simply "sensitive strain") and let $M_0$ be the sensitive strain's MIC. Let $\overline{M}$ be the MIC of the most-resistant strain and, for each $M > M_0$ let "strain $M$" denote all intermediate-resistant strains having MIC $M$. Let $X_0$ be the standard dose that would be prescribed absent resistance diagnosis, inducing recovery rate $\gamma_1 = \gamma(X_0; M_0)$ for those with sensitive infection and $\gamma(X_0;\ M) < \gamma_1$ for those with intermediate-resistant infection.

Now, suppose that POC-RD is available so that doctors can determine the extent of drug-1 resistance. Those found to be infected with intermediate-resistant bacteria could then be targeted with a higher dose, reducing or even reversing the treatment-survival advantage that these bacteria would otherwise normally enjoy under the standard dose. Whether such resistance-targeted dosing is enough to reverse the rise of intermediate resistance depends on how resistant bacteria have already become and how high the dose can be safely increased.

Suppose that the extent of resistance is sufficiently slight that the most-resistant strain can be cleared more quickly at the maximal safe dose (call it $\bar{X}$) as the sensitive strain is at the standard dose, i.e., $\gamma(\bar{X};\bar{M}) > \gamma_1$. POC-RD then enables doctors to hold all intermediate-resistant strains at a reproductive disadvantage relative to the sensitive strain, through a policy of (i) treating sensitive infections at the standard dose and (ii) treating intermediate-resistant infection at the maximal safe dose.[2] Under such targeted heightened dosing, the sensitive strain will have reproduction number $R_{0,0} = \frac{\beta}{\gamma_1}$ and each intermediate-resistant strain will have reproduction number $R_{0,M} = \frac{\beta}{\gamma(\bar{X};M)} < \frac{\beta}{\gamma(\bar{X};\bar{M})} < \frac{\beta}{\gamma_1}$, putting the sensitive strain at a reproductive advantage over all intermediate-resistant strains.

### Extension: Diagnostic delay

The analysis thus far assumes that the results of resistance diagnosis (RD) are available at the point of care, allowing resistant infections to be immediately treated and controlled as effectively as possible. Here we extend that baseline analysis to the case considered in the manuscript (MS), in which doctors learn the resistance profile of a patient's infection after delay $D \geq 0$ and can then adjust what antibiotic the patient is receiving and/or what transmission-control measures are put in place.

To keep equations as simple as possible, and to focus on the most challenging case, we will henceforth assume that isolation capacity is unavailable ($\epsilon = 0$) or, equivalently, that whatever resistant strains are currently in circulation are already so widespread that any attempt to isolate resistant carriers has negligible epidemiological impact.

**RD-targeted medical intervention**. When there is diagnostic delay, medical-care policies are more complex in that they specify how to treat and/or control infections initially at the point of care, as well as after RD results become available. For concreteness, we will restrict attention to policies in which (i) at the point of care, all patients are treated with drug 1 and no one is treated with drug 2; (ii) those found to have sensitive infection continue to receive drug-1 treatment; (iii) those found to have drug-1-resistant infection are put on drug-2 treatment and subjected to HTC; (iv) those found to have drug-2-

[2] In this context, even lower doses suffice to hold less-resistant strains at a reproductive disadvantage. For each MIC $M$, define $\underline{X}(M)$ so that $\gamma(\underline{X}(M);M) = \gamma_1$; in words, $\underline{X}(M)$ is the lowest dose that clears strain-$I_M$ infection at least as fast as the standard dose clears strain-$I_0$ infection. (Given the maintained assumption here that $\gamma(\bar{X};\bar{M}) > \gamma_1$, $\underline{X}(M) < \bar{X}$ and hence $\underline{X}(M)$ is a safe dose for each $M \leq \bar{M}$.) So long as each intermediate-resistant strain $I_M$ is targeted with a dose $X(M) > \underline{X}(M)$, the sensitive strain will enjoy a strict reproductive advantage.

resistant infection continue left on drug-1 treatment but subjected to HTC; and (v) those found to have pan-resistant infection are taken off antibiotic treatment and subjected to HTC.

*Discussion: Pre-diagnostic transmission control*. While waiting for the results of resistance diagnosis, doctors could order that infected hosts be subjected to HTC. Such an approach would reduce resistant-infection transmission but, since it also reduces sensitive-infection transmission, would have no overall effect on the resistant strain's *relative* reproductive fitness. It is therefore without loss for present purposes to restrict attention to policies that impose HTC only after a patient is confirmed as having resistant infection. That said, pre-diagnostic transmission-control efforts can be important in practice, especially when screening hosts who migrate into the host population. For instance, hospitals routinely isolate newly-admitted patients who are at risk of carrying carbapenem-resistant *Enterobacteriaceae* (CRE) in carriage, until they can be tested for CRE carriage. Such programs reduce the epidemiological impact of resistant in-migration, allowing hospitals to prevent CRE outbreaks (1, 2). See Part D for an extension allowing for host migration.

*Discussion: Diagnostic delay and followup care*. An important implicit assumption here is that treatment and transmission-control effort can be adjusted once resistance diagnosis is complete. However, in some cases, "the point of care" may be the only opportunity that doctors have to prescribe treatment or provide information to patients that can influence their behavior (and hence their subsequent transmissibility). If so, any test that takes longer than a patient remains at "the point of care" has the same effect on that patient's care as if resistance diagnosis were not available, i.e., as if delay $D = \infty$.

**Reproduction number and threshold fitness cost for each strain**

Let $R_{0,X}(D)$ denote strain X's reproduction number, where we add "$D$ notation" to emphasize that the formulas here depend on diagnostic delay $D \geq 0$. ($D = 0$ corresponds to the special case of POC-RD analyzed earlier. $D = \infty$ corresponds to the case of no RD.)

*Sensitive strain*. Strain 1 is unimpacted by RD, since all strain-1 infections continue to be treated with drug 1 and continue to be subjected to standard transmission control. Diagnostic delay therefore has no impact on strain 1's reproductive success:

$$R_{0,0}(D) = \frac{\beta}{\gamma_1} \text{ for all } D \geq 0$$

*Equation 2*

Drug-1-resistant strain. Absent RD, strain 1's reproduction number would be $R_{0,1}(\infty) = \frac{(1-f_1)\beta}{\gamma_0}$, where $f_1$ is the biological fitness cost of drug-1 resistance. The effect of RD is that, if a strain-1 infection lasts for length of time $D$, providers will switch to drug-2 treatment and heighten transmission control efforts upon detecting that the infection is resistant to drug 1, reducing the expected number of *subsequent* transmission events from $\frac{(1-f_1)\beta}{\gamma_0}$ to $\frac{Z_1(1-f_1)\beta}{\gamma_2}$. Since strain-1 infections clear at rate $\gamma_0$ during the "pre-diagnostic phase" until resistance diagnosis is complete, such infections last long enough to be detected with probability $e^{-D\gamma_0} < 1$. Overall, then, strain 1's reproduction number given diagnostic delay $D \geq 0$ is

$$R_{0,1}(D) = (1 - f_1)\beta\left(\frac{1}{\gamma_0} - e^{-D\gamma_0}\left(\frac{1}{\gamma_0} - \frac{Z_1}{\gamma_2}\right)\right)$$

*Equation 3*

Setting Eq. SI.3 equal to Eq. SI.2 and solving for $f_1$ yields the threshold fitness cost $f_1^*(D)$ for drug-1 resistance:

$$f_1^*(D) = 1 - \frac{1/\gamma_1}{\frac{1 - e^{-D\gamma_0}}{\gamma_0} + \frac{Z_1 e^{-D\gamma_0}}{\gamma_2}}$$

Figure 2 in the manuscript illustrates how $f_1^*(D)$ varies with diagnostic delay in a numerical example.

Drug-2-resistant and pan-resistant strains. Absent RD, strain 2 and strain 12 would have reproduction numbers $R_{0,2}(\infty) = \frac{(1-f_2)\beta}{\gamma_0}$ and $R_{0,12}(\infty) = \frac{(1-f_{12})\beta}{\gamma_0}$, respectively. If such infections last for length of time $D$, providers will heighten transmission control efforts ($HTC_2$ or $HTC_{12}$), reducing expected subsequent transmission from $\frac{\beta_2}{\gamma_1}$ to $\frac{Z_2\beta_2}{\gamma_1}$ for strain-2 infections or from $\frac{\beta_{12}}{\gamma_0}$ to $\frac{Z_{12}\beta_{12}}{\gamma_0}$ for strain-12 infections. Since strain-2 and strain-12 infections clear at respective rates $\gamma_1$ and $\gamma_0$ during the pre-diagnostic phase, such infections last long enough to be detected with respective probabilities $e^{-D\gamma_1}$ and $e^{-D\gamma_0}$. Overall, then, these strains reproduction numbers are

$$R_{0,2}(D) = \frac{(1 - f_2)\beta}{\gamma_1}\left(1 - e^{-D\gamma_1}(1 - Z_2)\right)$$

*Equation 4*

$$R_{0,12}(D) = \frac{(1 - f_{12})\beta}{\gamma_0}\left(1 - e^{-D\gamma_0}(1 - Z_{12})\right)$$

*Equation 5*

Observe that $R_{0,2}(D) < R_{0,0}(D)$ for all $D < \infty$; so, selection against drug-2 resistance can be maintained no matter what the diagnostic delay, even if there are no biological fitness costs associated with drug-2 resistance. On the other hand, so long as $Z_{12} > \frac{\gamma_0}{\gamma_1}$, meaning that $HTC_{12}$ is less effective than drug-1 treatment at preventing transmission, then $R_{0,12}(D) > R_{0,0}(D)$ for all $D \geq 0$. We conclude that, even with zero diagnostic delay, selection against pan resistance requires *either* that there be sufficiently large biological fitness benefits associated with pan resistance *or* that sufficiently effective transmission-control options (i.e., isolation) are available that can be targeted against the pan-resistant strain. Setting Eq. SI.5 equal to Eq. SI.2 and solving for $f_{12}$ yields threshold fitness cost ${f_{12}}^*(D)$ for pan resistance:

$${f_{12}}^*(D) = 1 - \frac{\gamma_0/\gamma_1}{1 - (1 - Z_{12})e^{-D\gamma_0}}$$

*Equation 6*

Figure 2 in the manuscript illustrates how ${f_{12}}^*(D)$ varies with diagnostic delay in a numerical example.

### Extension: Heightened discovery

Here we further extend the benchmark model to allow for both heightened discovery (causing resistant-pathogen carriers to come to medical attention more quickly) and diagnostic delay. Suppose that hosts infected by strain X=0,1,2,12 come to medical attention at rate $r_X$, after which their infections are treated and controlled as in the previous section, informed by RD with diagnostic delay $D \geq 0$. We assume that there is no reason why resistant infections are inherently harder to discover, i.e., $r_0 \leq \max\{r_1, r_2, r_{12}\}$, but allow for the possibility of accelerated discovery of resistant infections, as discussed in Part A. If $r_X > r_0$, we say that there is “heightened discovery of strain $X$ ($HD_X$)”.

<u>*Discussion: heightened discovery.*</u> One can interpret $r_0$ as the rate at which infected hosts seek out medical care and $r_X - r_0$ as the rate at which hosts with strain-$X$ infection are “discovered” and brought under medical care before they would have otherwise sought out care. There are many ways in which to speed discovery of resistant infection. Some examples: *Contact tracing*: When a patient is found with

resistant infection, close contacts (at home, in school or the workplace, etc.) could be pre-emptively tested and brought under medical care if found to be harboring resistant bacteria. *Contact notification*: If testing a patients' epidemiological contacts is not feasible, it may still be possible to notify them that they may have been exposed to resistant infection and encourage them to seek out care as soon as they begin to experience symptoms.[3] *Routine screening for resistance:* Hosts harboring resistant bacteria sometimes seek out medical care for conditions unrelated to those bacteria. Routine testing at those points of care would allow doctors to identify when hosts are harboring resistant bacteria. So long as doctors are more likely to act on this knowledge when resistant bacteria are found, such routine screening will increase resistant-infection discovery more than it increases sensitive-infection discovery.

**Reproduction number and threshold fitness cost for each strain**

Let $R_{0,X}(r_X, D)$ denote strain X's reproduction number, where now we add "$r_X$ notation" to emphasize that the formulas here depend on the rate at which strain-X infections are discovered as well as diagnostic delay.

<u>*Sensitive strain:*</u> Absent infection discovery, strain-0 infections would never be treated and strain 0's reproduction number would be $R_{0,0}(\infty, D) = \frac{\beta}{\gamma_0}$. If a strain-0 infection lasts long enough to be discovered, providers will initiate drug-1 treatment, reducing the expected number of subsequent transmission events from $\frac{\beta}{\gamma_0}$ to $\frac{\beta}{\gamma_1}$. Since strain-0 infections clear at rate $\gamma_0$ and are discovered at rate $r_0$ during the "pre-discovery phase," hosts infected by strain 0 are discovered with probability $\frac{r_0}{r_0+\gamma_0}$. Overall, then, strain 1's reproduction number given discovery rate $r_0$ and diagnostic delay $D \geq 0$ is $R_{0,0}(r_0, D) = \beta\left(\frac{1}{\gamma_0} - \frac{r_0}{r_0+\gamma_0}\left(\frac{1}{\gamma_0} - \frac{1}{\gamma_1}\right)\right)$. Re-arranging terms yields an equivalent alternative formulation, which may be more intuitive for some readers:

$$R_{0,0}(r_0, D) = \beta\left(\frac{1}{r_0 + \gamma_0} + \frac{r_0}{r_0 + \gamma_0} \times \frac{1}{\gamma_1}\right)$$

*Equation 7*

[3] Some contacts of a patient with resistant infection will develop sensitive infection; so, contact notification will tend to increase both sensitive- and resistant-infection discovery. However, because the contacts of a patient with resistant infection are more likely to have resistant infection than the general population, the overall effect is to increase resistant-infection discovery more than sensitive-infection discovery.

To understand this second formulation, note that sensitive infections remain in the pre-discovery phase for average length of time $\frac{1}{r_0+\gamma_0}$, get discovered with probability $\frac{r_0}{r_0+\gamma_0}$ and, if discovered, persist on average for an additional length of time $R_{0,0}(0,D)=\frac{1}{\gamma_1}$ (Eqn SI.2). We use this second formulation in the analysis to follow, when deriving resistant strains' reproduction numbers.

*Drug-1-resistant strain:* During the pre-discovery phase, strain-1 infections are discovered at rate $r_1$ and clear at rate $\gamma_0$. The pre-discovery period for strain-1 infections therefore lasts on average for length of time $\frac{1}{r_1+\gamma_0}$ and such infections come to medical attention with probability $\frac{r_1}{r_1+\gamma_0}$, after which the expected number of subsequent transmission events is $R_{0,1}(0,D)=(1-f_1)\beta\left(\frac{1}{\gamma_0}-e^{-D\gamma_0}\left(\frac{1}{\gamma_0}-\frac{Z_1}{\gamma_2}\right)\right)$ (Eqn SI.3), the same *as if* the infection had been immediately discovered with diagnostic delay $D$. Strain 1's reproduction number is therefore

$$R_{0,1}(r_1,D)=(1-f_1)\beta\left(\frac{1}{r_1+\gamma_0}+\frac{r_1}{r_1+\gamma_0}\times\left(\frac{1}{\gamma_0}-e^{-D\gamma_0}\left(\frac{1}{\gamma_0}-\frac{Z_1}{\gamma_2}\right)\right)\right)$$

*Equation 8*

Setting Eq. SI.7 equal to Eq. SI.8 and solving for $f_1$ yields the threshold fitness cost $f_1{}^*(r_0,r_1,D)$ for drug-1 resistance in this context:

$$f_1^*(r_0,r_1,D)=1-\frac{\frac{1}{r_0+\gamma_0}\times\left(1+\frac{r_0}{\gamma_1}\right)}{\frac{1}{r_1+\gamma_0}\times\left(1+r_1\left(\frac{1}{\gamma_0}-e^{-D\gamma_0}\left(\frac{1}{\gamma_0}-\frac{Z_1}{\gamma_2}\right)\right)\right)}$$

*Equation 9*

Note that $f_1^*(r_0,r_1,D)<0$—meaning that net selection against drug-1 resistance can be maintained even if there are no biological fitness costs—so long as (i) drug 2 is equally effective as drug 1 ($\gamma_2=\gamma_1$), (ii) diagnostic delay is negligible (so that $e^{-D\gamma_0}\approx 1$), and (iii) *either* $HTC_1$ is at least somewhat effective ($Z_1<1$) *or* $HD_1$ is at least somewhat effective ($r_1>r_0$) .

*Drug-2-resistant strain:* Because strain-2 infections clear at rate $\gamma_0$ and are discovered at rate $r_2$, the pre-discovery period lasts for average time $\frac{1}{r_2+\gamma_0}$ and such infections come to medical attention with probability $\frac{r_2}{r_2+\gamma_0}$. After coming to medical attention, expected subsequent transmission is then $R_{0,2}(D) = \frac{(1-f_2)\beta}{\gamma_1}\left(1 - e^{-D\gamma_1}(1 - Z_2)\right)$ (Eqn. SI.4). The reproduction number for strain 2 is therefore

$$R_{0,2}(r_2, D) = (1 - f_2)\beta\left(\frac{1}{r_2+\gamma_0} + \frac{r_2}{r_2+\gamma_0} \times \frac{1}{\gamma_1}\left(1 - e^{-D\gamma_1}(1 - Z_2)\right)\right)$$

*Equation 10*

Setting Eq. SI.7 equal to Eq. SI.10 and solving for $f_2$ yields the threshold fitness cost $f_2{}^*(r_0, r_2, D)$ for drug-2 resistance:

$$f_2^*(r_0, r_2, D) = 1 - \frac{\frac{1}{r_0+\gamma_0} \times \left(1 + \frac{r_0}{\gamma_1}\right)}{\frac{1}{r_2+\gamma_0} \times \left(1 + \frac{r_2}{\gamma_1}\left(1 - e^{-D\gamma_1}(1 - Z_2)\right)\right)}$$

*Equation 11*

Note that the threshold fitness cost for drug-2 resistance $f_2^*(r_1, D) \leq 0$ even if $HTC_2$ and $HD_2$ are completely ineffective ($Z_2 = 1$ and $r_2 = r_0$) and no matter what the diagnostic delay.

*<u>Pan-resistant strains</u>:* Because strain-12 infections clear at rate $\gamma_0$ and are discovered at rate $r_{12}$, the pre-discovery period lasts for average time $\frac{1}{r_{12}+\gamma_0}$ and such infections come to medical attention with probability $\frac{r_{12}}{r_{12}+\gamma_0}$, after which subsequent transmission is $R_{0,12}(D) = \frac{(1-f_{12})\beta}{\gamma_0}\left(1 - e^{-D\gamma_0}(1 - Z_{12})\right)$ (Eqn. SI.5). The reproduction number for strain 12 is therefore

$$R_{0,12}(r_{12}, D) = (1 - f_{12})\beta\left(\frac{1}{r_{12}+\gamma_0} + \frac{r_{12}}{r_{12}+\gamma_0} \times \frac{1}{\gamma_0}\left(1 - e^{-D\gamma_0}(1 - Z_{12})\right)\right)$$

*Equation 12*

Setting Eq. SI.7 equal to Eq. SI.12 and solving for $f_{12}$ yields the threshold fitness cost $f_{12}{}^*(r_0, r_{12}, D)$ for pan resistance:

$$f_{12}^{*}(r_0, r_{12}, D) = 1 - \frac{\frac{1}{r_0+\gamma_0} \times \left(1 + \frac{r_0}{\gamma_1}\right)}{\frac{1}{r_{12}+\gamma_0} \times \left(1 + \frac{r_{12}}{\gamma_0}\left(1 - e^{-D\gamma_0}(1 - Z_{12})\right)\right)}$$

*Equation 13*

To explore the implications of Eqn SI.12, consider the relatively simple case without any diagnostic delay ($D = 0$). In this case, the threshold fitness cost of pan resistance $f_{12}^{*}(r_0, r_{12}, 0) = 1 - \frac{\frac{1}{r_0+\gamma_0} \times \left(1+\frac{r_0}{\gamma_1}\right)}{\frac{1}{r_{12}+\gamma_0} \times \left(1+\frac{r_{12}Z_{12}}{\gamma_0}\right)}$.

Note that, if there are no biological fitness costs, it may still be possible to maintain a net selection against the pan resistant strain, but only if

$$\frac{1}{r_0+\gamma_0} \times \left(1 + \frac{r_0}{\gamma_1}\right) > \frac{1}{r_{12}+\gamma_0} \times \left(1 + \frac{r_{12}Z_{12}}{\gamma_0}\right)$$

*Equation 14*

If pan-sensitive and pan-resistant infections are discovered equally rapidly ($r_0 = r_{12}$), then the inequality in Eqn SI.14 only holds if $Z_{12} < \frac{\gamma_0}{\gamma_1}$, meaning that $\text{HTC}_{12}$ measures are even more effective than drug-1 treatment at preventing transmission. But what if $\text{HTC}_{12}$ measures are only moderately effective ($1 > Z_{12} > \frac{\gamma_0}{\gamma_1}$)? How effective must $\text{HD}_{12}$ measures be in order to maintain a net selection against the pan-resistant strain?

First, we observe that even the most intensive possible HD efforts may not be enough on their own to select against the pan-resistant strain. To see why, note that the right-hand-side (RHS) of Eqn. SI.14 converges to $\frac{Z_{12}}{\gamma_0}$ as $r_{12} \to \infty$, while the left-hand-side (LHS) converges to $\frac{1}{\gamma_1}$ as $r_0 \to \infty$. Thus, so long as the baseline rate $r_0$ at which pan-sensitive infections is sufficiently high, even *immediate* discovery of pan-resistant infections is insufficient to hold the pan-resistant strain at a disadvantage.

On the other hand, in cases when pan-sensitive infections are unlikely to be discovered ($r_0 \approx 0$), sufficiently-effective heightened discovery of pan-resistant infection—coupled with moderately effective HTC measures that can be applied, once such infections are discovered—will always suffice to select against the pan-resistant strain. To see why, consider the limit in which $r_0 \to 0$ and $r_{12} \to \infty$. In this limit, the LHS of Eqn SI.14 converges to $\frac{1}{\gamma_0}$ while the RHS converges to $\frac{Z_{12}}{\gamma_0} < \frac{1}{\gamma_0}$, implying that $\lim_{r_0 \to 0, r_{12} \to \infty} f_{12}^*(r_0, r_{12}, 0) < 0$, meaning that net selection against the pan-resistant strain can be maintained even when there are no biological fitness costs associated with pan resistance.[4]

### Extension: Sustainable antibiotic use

Our analysis so far in Part B has focused on whether resistant strains of an obligate pathogen can be held at a reproductive disadvantage *even if* there are no biological fitness costs associated with resistance. We have identified several contexts in which this may be possible. For instance, *selection against the drug-1-resistant strain* can be achieved without withholding treatment from anyone so long as an equally-effective alternative treatment (drug 2) is available to which the drug-1-resistant strain remains susceptible, by using POC-RD to identify strain-1 infections and then treating them with drug 2 and deploying heightened transmission controls ($\text{HTC}_1$). Similarly, *selection against the pan-resistant strain* can be achieved without withholding treatment so long as infections are rarely discovered, by

[4] Similar results can be easily established in the more complex case with diagnostic delay; in particular, $\lim_{r_0 \to 0, r_{12} \to \infty} f_{12}^*(r_0, r_{12}, D) < 0$ for all $D \geq 0$.

deploying heightened discovery efforts ($HD_{12}$) to increase the likelihood that pan-resistant infections are brought to medical attention and identified by RD, and then deploying $HTC_{12}$. In these cases, all patients can be sustainably treated with the most effective antibiotic for their infection without promoting the rise of antibiotic resistance, even if there are no biological fitness costs.

On the other hand, we have also identified scenarios in which RD-informed discovery, treatment, and control is not sufficient to maintain a net selection against some resistant strains. For instance, *selection against the pan-resistant strain* cannot be achieved while also treating all patients, so long as infections are very likely to be discovered (limiting the impact of $HD_{12}$) and transmission-control options are only somewhat effective (limiting the impact of $HTC_{12}$). In these more challenging cases, if there are no biological fitness costs, treating all patients with the most effective antibiotic for their infection will lead resistant strains to grow more prevalent over time, undermining our ability to treat infections effectively in the future.

In these cases when all patients cannot be sustainably treated, other investments *beyond resistance diagnostics* are needed to "change the game" and enable circulating resistant strains to be controlled. In this section, we provide a quantitative measure of the remaining challenge ("maximal sustainable antibiotic use," denoted by $\alpha^*$) to assess, for each pathogen of interest, the comparative benefit of (say) investing in developing a new antibiotic versus investing to enhance public-health capabilities to strengthen RD-informed HTC and/or HD options.

**Maximal sustainable antibiotic use**

Suppose that all resistant infections are treated and controlled as effectively as possible but that only fraction $\alpha \in [0,1]$ of those with pan-sensitive infection receive antibiotic treatment. Moreover, to simplify equations and focus on the most challenging case, suppose that there are no biological fitness costs associated with resistance. Let $\alpha^*$ denote the largest fraction of pan-sensitive infections that can be treated with drug 1 while maintaining a net selection against all resistant strains currently in circulation. We refer to $\alpha^*$ as "maximal sustainable antibiotic use".

Withholding treatment from fraction $(1-\alpha)$ of those with pan-sensitive infection increases strain 0's reproduction number but has no effect on the reproduction numbers of the resistant strains (given by Eqns. SI.8, SI.10, and SI.12). Once brought to medical attention, strain-0 infections that are treated last on average for duration $\frac{1}{\gamma_1}$, while those that are left untreated last on average for $\frac{1}{\gamma_0}$. Overall, then, the expected duration of strain-0 infections after discovery is $\frac{\alpha}{\gamma_1} + \frac{1-\alpha}{\gamma_0}$ and strain 0's reproduction number can be derived by appropriately modifying Eqn SI.7:

$$R_{0,0}(r_0, D, \alpha) = \beta\left(\frac{1}{r_0 + \gamma_0} + \frac{r_0}{r_0 + \gamma_0} \times \left(\frac{\alpha}{\gamma_1} + \frac{1-\alpha}{\gamma_0}\right)\right)$$

*Equation 15*

For each resistant strain X=1,2,12, let $\alpha_X^*$ denote the maximal value of $\alpha$ given which $R_{0,0}(r_0, D, \alpha) \geq R_{0,X}(r_X, D)$. $\alpha_1^*$, $\alpha_2^*$, and $\alpha_{12}^*$ can be derived by setting Eqn SI.15 equal to Eqns SI.8, SI.10, and SI.12 (setting $f_1 = f_2 = f_{12} = 0$ because of the maintained assumption that there are no biological fitness costs), respectively, and solving for $\alpha$. (We omit details because the resulting formulas are complex and do not provide additional insight.) If drug-2 resistance has not yet emerged, so that only strain 1 is in circulation, then maximal sustainable antibiotic use $\alpha^* = \alpha_1^*$; otherwise, if all three resistant strains are in circulation, then $\alpha^* = \min\{\alpha_1^*, \alpha_2^*, \alpha_{12}^*\}$.

Figure SI.4 illustrates how $\alpha^*$ varies with diagnostic delay $D$ in a simple numerical example in which there are no biological fitness costs and no scope for heightened discovery because all infections are immediately brought to medical attention. Three features of Figure SI.4 are worth highlighting. First, if RD is unavailable, no amount of drug-1 use is sustainable once the drug-1-resistant strain is in circulation, whether or not the pan-resistant strain is also in circulation. Second, once the pan-resistant strain is in circulation, RD-informed interventions allow up to 20% of those with pan-sensitive infection to be sustainably treated, but it is not possible to sustainably treat all patients. On the other hand, if the pan-resistant strain is not yet in circulation, all patients can be sustainably treated so long as resistance-diagnostic delay is less than a critical threshold $D^*$ (about one day in the numerical example).

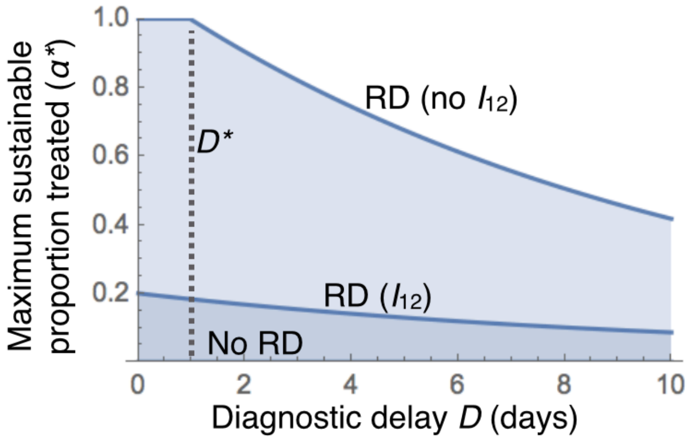


**Figure SI.4. Rapid resistance diagnostics enable conditional treatment and infection control strategies that can select against resistance for obligate pathogens even with widespread antibiotic use**. The maximal proportion of sensitive infections that can be treated without causing an increase in resistance ($\alpha^*$) is plotted against diagnostic delay $D$, assuming that all infections are immediately discovered and there are no biological fitness costs. The dashed vertical line indicates the longest diagnostic delay ($D^*$) given which there is selection against drug-1 resistance while treating all cases. Three scenarios are shown: resistance diagnostics (RD) not available (No RD), for which $\alpha^* = 0$; RD available with delay and pan-resistance not yet emerged (RD, no $I_{12}$); RD available with delay and pan-resistance widespread (RD, $I_{12}$). Parameters (rates per day): $\gamma_0^I = 0.1$, $\gamma_1^I = \gamma_2^I = 0.2$, $\beta^I = 0.2$.

### Part C. An opportunistic pathogen

This section extends the obligate-pathogen analysis of Part B to a setting in which the pathogen of interest ("Pathogen 1") is an opportunistic pathogen that dwells in an asymptomatic "carriage state" before progressing to cause infection and faces occasional episodes of antibiotic exposure when hosts receive antibiotic treatment for infections caused by other pathogens. If these other infections are treated with drug 1 and Pathogen 1 is sufficiently unlikely to cause infection, the resulting *untargeted bystander exposure* can create an irreversible selective pressure favoring drug-1-resistant and totally-resistant strains of Pathogen 1 (Proposition C.2)—unless these resistant strains of Pathogen 1 can be discovered and controlled before they cause infection (Proposition C.3).

**Model: opportunistic pathogen (Pathogen 1 or P1).** As in Part B, there are two available antibiotics (drug 1 and drug 2) and up to four strains of Pathogen 1 in circulation: $I_0$, sensitive to both drugs; $I_1$, resistant to drug 1 but sensitive to drug 2 ("drug-1 resistant"); $I_2$, resistant to drug 2 but sensitive to drug 1 ("drug-2 resistant"); and $I_{12}$, resistant to both drugs ("totally resistant"). Pathogen 1 circulates according to a standard susceptible-carriage-infected (SCIS) model, potentially with transmission both from carriage and during infection. *Transmission:* The baseline transmission rate during infection is $\beta^I \geq 0$ (same for all strains) but, for each resistant strain $I_X$ (X=1,2,12), heightened transmission control measures ("infection HTC") are available that can reduce strain-$I_X$ transmission during infection to $Z_X^I \beta^I$, for some $0 \leq Z_X^I \leq 1$. The baseline transmission rate during carriage is $\beta^C \geq 0$ (same for all strains). In an extension, we consider the impact of "carriage HTC" measures that can reduce strain-$I_X$ transmission during carriage. For the moment, however, assume that there is no way to control the pathogen prior to infection. *Progression to infection*: While in carriage, Pathogen 1 progresses to cause infection at rate $d^C > 0$. (If $d^C = \infty$, Pathogen 1 spends no time in carriage and is therefore effectively an obligate pathogen as in Part B.) For simplicity, we assume that infections are immediately discovered. *Clearance during infection:* Same as in Part B, with the same notation. Infected hosts recover at rate $\gamma_0 > 0$ if left untreated or ineffectively treated, at rate $\gamma_1 > \gamma_0$ if effectively treated with drug 1, and at rate $\gamma_2 > \gamma_0$ if effectively treated with drug 2. *Clearance during carriage:* Pathogen 1 clearance from carriage can arise for two sorts of reasons: (i) Immune response, microbiome interactions, etc, that clear P1 at baseline rate $\gamma^C > 0$ (same for all strains). (ii) Bystander exposure (BE) events arising at rate $\eta > 0$ as hosts are treated for other infections and each clearing P1 from carriage with probability $p_x^{BE}$ when drug x is used and a drug-x-sensitive strain of P1 is in carriage. For convenience, let $\phi_x^C = \eta p_1^{BE}$, the overall rate at which drug-x-sensitive P1 is cleared from carriage due to bystander exposure to drug x. (Overall, the sensitive strain is cleared due to BE at rate $\phi_1^C + \phi_2^C$, the drug-1-resistant strain is cleared at rate $\phi_2^C$, and the drug-2-resistant strain is cleared at rate $\phi_1^C$.)

_Discussion: other infections and bystander exposure (BE) in carriage._ Each host contracts an infection caused by another pathogen (Pathogen 2 or P2) at fixed rate $\eta > 0$. For analytical simplicity, we assume that P2-infections are short-lived relative to the average length of time that Pathogen 1 spends in carriage, modeling such infections—and resulting antibiotic exposure—as occurring at particular moments in time.[5] Also for simplicity, we assume: (i) there are no resistance concerns with Pathogen 2, i.e., all P2-infections are sensitive to both drugs and (ii) drug 1 is the preferred treatment for P2-infection but, if drug-1-resistant Pathogen 1 were found in carriage, doctors would prefer to prescribe drug 2.

_Discussion: co-colonization dynamics_. Implicit in our modeling approach is a simplifying assumption that Pathogen 1's colonization and infection dynamics are the same whether or not Pathogen 2 is also present in the host, and vice versa. In future work, it would be interesting to extend our analysis to settings with richer co-colonization dynamics. For instance, suppose that P2 is an opportunistic pathogen that is more likely to progress to cause infection when P1 is also in carriage. If P1 is more widespread in carriage, more hosts will contract P2 infection, creating more opportunities to conduct carriage RD and hence to target resistant P1-strains with interventions aimed at reducing transmission from carriage. On the other hand, if P1 is rare, fewer hosts will contract P2 infection, creating fewer opportunities to clear resistant P1 strains before they can progress to cause infection.

#### Baseline Case: All infections treated with drug 1 and controlled equally

We begin by deriving the reproduction number of each Pathogen 1 strain in the relatively simple baseline case in which neither infection POC-RD nor carriage RD is available. In this case, all infections are treated with drug 1 and Pathogen 1 is never subjected to heightened transmission control during infection or during carriage. Because by assumption there are no biological fitness costs associated with resistance, the drug-1-resistant strain $I_1$ and the totally-resistant strain $I_{12}$ obviously enjoy a

[5] The analysis can be extended to a more realistic setting in which treated P2-infections clear at rate $\gamma^{P2} > 0$ and hence last for average length of time $1/\gamma^{P2}$. In particular, suppose that drug-1 exposure clears P1 from carriage at rate $\gamma_1^{BE} > 0$. Given baseline carriage clearance rate $\gamma^C > 0$ and disease-development rate $d^C > 0$, each episode of drug-1 exposure will clear P1 from carriage with probability $\frac{\gamma_1^{BE}}{\gamma^C + \gamma_1^{BE} + d^C} > 0$, and likewise for drug-2 exposure.

reproductive advantage over the sensitive strain $I_0$. Even so, it is useful to derive the strains' reproduction numbers $R_{0,0}$, $R_{0,1}$ and $R_{0,12}$ in this baseline case, since the method of derivation can be adapted and extended to other more interesting cases.

**Proposition C.1**: *Suppose that resistance diagnosis (RD) is not available. The drug-1 resistant strain $I_1$ and pan-resistant strain $I_{12}$ enjoy an unambiguous reproductive advantage over the sensitive strain $I_0$, with $R_{0,12} = R_{0,1} > R_{0,0}$.*

The rest of this section verifies Proposition C.1.

**Reproduction number analysis.** Upon entering a new host, P1 begins in a "carriage phase" until either being cleared from carriage or progressing to an "infection phase". For each resistance profile $X \in \{0,1,2,12\}$, let $L_X^C$ be strain $I_X$'s average length of time spent in the carriage phase, let $P_X^{C\to I}$ be the probability that strain $I_X$ progresses to cause infection, and let $\Pi_X^I$ be the expected number of transmission events (when the pathogen is transmitted to a new host, including cases when the host is not susceptible to colonization) during strain-$I_X$ infection. Because the pathogen is transmitted at rate $\beta^C$ while in carriage, strain $I_X$'s reproduction number is

$$R_{0,X} = \beta^C L_X^C + P_X^{C\to I} \Pi_X^I$$

*Equation C.1*

<u>Reproductive success during infection</u>: The terms $\Pi_X^I$ were derived in Part B. To summarize: Since infection POC-RD is not available, all P1-infections are treated with drug 1 and subjected to standard transmission control (STC); so,

$$\Pi_0^I = \Pi_2^I = \beta^I \tau_1 \text{ and } \Pi_1^I = \Pi_{12}^I = \beta^I \tau_0$$

*Equation C.2*

<u>*Time spent in carriage*</u>**.** There are four ways in which the carriage phase may end: progression to infection (rate $d^C$ for all strains); baseline clearance (rate $\gamma^C$ for all strains); clearance due to incidental drug-1 exposure (rate $\phi_1^C$ for strains $I_0, I_2$, rate 0 for strains $I_1, I_{12}$); or clearance due to incidental drug-2 exposure (rate $\phi_2^C$ for strains $I_0, I_1$, rate 0 for strains $I_2, I_{12}$). Overall, then, the carriage phase ends at rate $d^C + \gamma^C + \phi_1^C + \phi_2^C$ for strain $I_0$, at rate $\gamma^C + d^C + \phi_2^C$ for strain $I_1$, at rate $d^C + \gamma^C + \phi_1^C$ for strain $I_2$, and at rate $\gamma^C + d^C$ for strain $I_{12}$. The expected length of time that each strain $I_X$ spends in the carriage phase is therefore

$$L_X^C = \frac{1}{d^C + \gamma^C + \sum_{x \notin X} \phi_x^C}$$

*Equation C.3*

<u>*Likelihood of progressing to infection*</u>. Each strain $I_X$ progresses to infection at rate $d^C$ and is cleared from carriage at rate $\gamma^C + \sum_{x \notin X} \phi_x^C$. The likelihood that strain $I_X$ progresses to infection before being cleared is therefore

$$P_X^{C \to I} = P_2^{C \to I} = \frac{d^C}{d^C + \gamma^C + \sum_{x \notin X} \phi_x^C}$$

*Equation C.4*

Combining Eqs. C.1-C.4, we find each strain's Baseline Reproduction Number:

$$R_{0,0} = \frac{\beta^C + d^C \beta^I / \gamma_1}{d^C + \gamma^C + \phi_1^C + \phi_2^C}, \; R_{0,1} = \frac{\beta^C + d^C \beta^I / \gamma_0}{d^C + \gamma^C + \phi_2^C}, \; R_{0,2} = \frac{\beta^C + d^C \beta^I / \gamma_1}{d^C + \gamma^C + \phi_1^C}, \; R_{0,12} = \frac{\beta^C + d^C \beta^I / \gamma_0}{d^C + \gamma^C}$$

*Equation C.5*

The fact that $R_{0,0} < R_{0,1}$ follows immediately from the fact that drug-1 treatment can clear the sensitive strain during infection ($\gamma_1 > \gamma_0$) and that drug-1 exposure can clear the sensitive strain in carriage ($\phi_1^C > 0$).

#### Impact of Introducing Infection POC-RD

**Proposition C.2**: *Suppose that infection POC-RD is available. So long as the frequency of Pathogen 2 infections* $\eta > \eta^* = \frac{\beta^I (d^C + \gamma^C) d^C / \gamma_1}{\beta^C p_1^{AE}}$*, the drug-1-resistant strain* $I_1$ *of P1 will remain at a reproductive advantage over the sensitive strain even if drug-1-resistant P1-infections can be perfectly quarantined.*

The rest of this section verifies Proposition C.1, by investigating how the availability of infection POC-RD changes how resistance strains can be treated and controlled.

<u>*Sensitive strain*</u>: Strain-$I_0$ infections receive the same (drug-1) treatment and the same (standard) transmission control. Strain $I_0$'s reproduction number is therefore also unchanged: $R_{0,0} = \frac{\beta^C + d^C \beta^I / \gamma_1}{d^C + \gamma^C + \phi_1^C + \phi_2^C}$. Recall that, by definition, $\phi_x^C = \eta p_x^{AE}$ for each drug *x=1,2*.)

*Drug-1-resistant strain*: Strain-$I_1$ infections now receive effective antibiotic treatment (with drug 2) as well as infection HTC, lowering strain $I_1$'s reproduction number to $R_{0,1} = \frac{\beta^C + d^C Z_1^I \beta^I/\gamma_2}{d^C+\gamma^C+\phi_2^C}$.

*Drug-2-resistant strain*: Strain-$I_2$ infections receive the same treatment (with drug 1) but are now subjected to infection HTC, lowering strain $I_2$'s reproduction number to $R_{0,2} = \frac{\beta^C + d^C Z_2^I \beta^I/\gamma_1}{d^C+\gamma^C+\phi_1^C}$.

*Totally-resistant strain*: Strain-$I_{12}$ infections remain untreatable, but now can at least be subjected to infection HTC, lowering strain $I_1$'s reproduction number to $R_{0,12} = \frac{\beta^C + d^C Z_{12}^I \beta^I/\gamma_0}{d^C+\gamma^C}$.

To verify the claim in Proposition C.2, suppose that quarantine is available to perfectly isolate those with drug-1-resistant infection, i.e., $Z_1^I = \infty$. The reproduction-number formula for strain $I_1$ then reduces to $R_{0,1} = \frac{\beta^C}{d^C+\gamma^C+\phi_1^C}$. Note that $R_{0,0} \geq R_{0,1}$ if and only if $\frac{\beta^C + d^C\beta^I/\gamma_1}{\beta^C} \geq \frac{d^C+\gamma^C+\phi_1^C}{d^C+\gamma^C}$, which simplifies to $\frac{d^C\beta^I/\gamma_1}{\beta^C} \geq \frac{\phi_1^C}{d^C+\gamma^C}$ or, equivalently, $\eta \leq \eta^* = \frac{\beta^I(d^C+\gamma^C)d^C/\gamma_1}{\beta^C p_1^{AE}}$.

In Fig SI.5, we examine our SCIS model with parameters illustrative for the pneumococcus (as discussed in the MS, details on the parametrization in SI.G), to determine the maximal sustainable antibiotic use[6] given POC-RD for a bacterium with substantial bystander exposure (Fig 3 in the MS). Even in the absence of pan-resistant strains, the long-carriage duration of most pneumococcal serotypes (median serotype-specific carriage duration is approximately 10 weeks) ensures that bystander selection dominates the relatively weak impact of POC-RD during rare and brief infection events (median infection duration in absence of treatment is 8 days).

[6] If fraction $1-\alpha$ of pan-sensitive infections are left untreated, strain 1's reproduction number is $R_{0,0}(\alpha) = \frac{\beta^C + d^C\beta^I\left(\frac{\alpha}{\gamma_1}+\frac{1-\alpha}{\gamma_0}\right)}{d^C+\gamma^C+\phi_1^C+\phi_2^C}$. Maximal sustainable antibiotic use $\alpha^*$ here is the highest given which $R_{0,0}(\alpha) \geq R_{0,1}$.

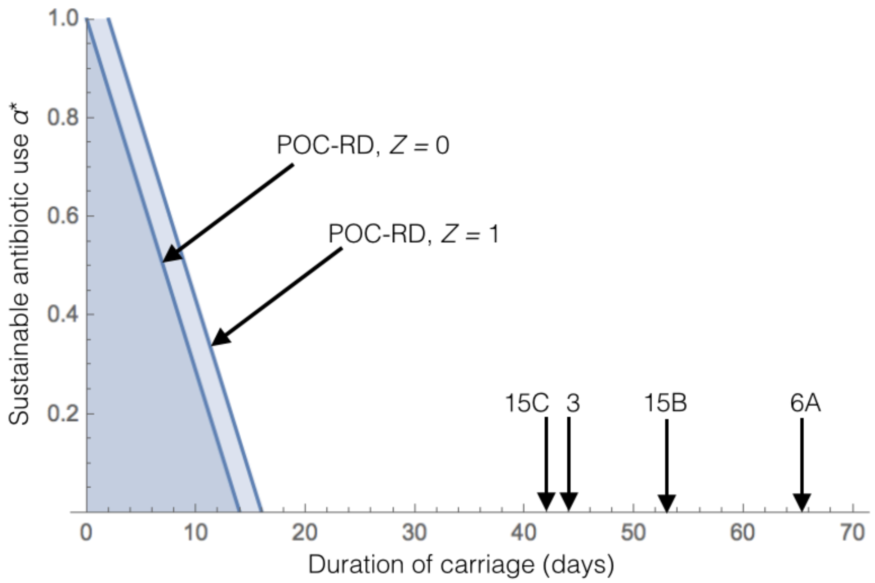


**Figure SI.5. POC-RD alone cannot reverse selection on pneumococcal resistance, due to long carriage times**. The maximal proportion of sensitive infections that can be treated without causing an increase in drug 1 resistance ($\alpha^*$) is plotted against the expected duration of carriage. Two POC-RD scenarios are shown: with ($Z_1^I = 0$) and without ($Z_1^I = 1$) transmission control. Vertical arrows represent pneumococcal serotypes with below-average carriage duration (see main text Fig. 4 for broader range of serotype carriage durations). The remaining parameters (rates per day) are $d$ = 0.001, $\phi_1^C$ = 5x10$^{-4}$, $\phi_2^C$ = 0, $\gamma_1^I = \gamma_2^I$ = 1, $\gamma_0^I$ = 0.125. We make the simplifying assumption that baseline carriage and infection transmission rates are identical ($\beta^C = \beta^I = \beta$) given which $\alpha^*$ does not depend on $\beta$. Details on parameterization are in SI.G.

#### Extension: Discovery and Control of Resistant Strains while in Carriage

This section provides an extension of the previous opportunistic-pathogen analysis, allowing for the possibility that it may be possible to discover and control Pathogen 1 strains even before they cause any symptoms. The analysis here provides the basis for Fig 5 in the manuscript.

An implicit assumption of the analysis thus far is that there is no way to treat and/or control the spread of resistant opportunistic pathogens until after they have progressed to cause symptomatic infection. But what if resistant strains of an opportunistic pathogen could be discovered while in carriage? Once discovered, hosts carrying a resistant strain could then be subjected to heightened transmission control measures, thereby reducing the resistant strain's transmissibility prior to infection. For example, in the South Swedish Pneumococcal Intervention Project (SSPIP), public-health officials tested all preschool-age children in the same daycare class whenever any of them was diagnosed with penicillin-resistant pneumococcal (PRP) infection, and required that any child found to have PRP in carriage remain home until they could be verified as PRP-free. (SSPIP is discussed in more detail in the main manuscript.)

**Extensions to the model**. To explore the effect of public-health interventions such as SSPIP, we add two features to the opportunistic-pathogen model. *Discovery during carriage:* For each resistance profile X = 0,1,2,12, let $r_X \geq 0$ be the rate at which strain $I_X$ is discovered while in carriage. *Transmission control during carriage*: Once discovered in carriage, strain $I_X$ can be subjected to carriage HTC measures that decrease transmission from carriage by a factor of $Z_X^C \leq 1$. The baseline model corresponds to the special case in which $r_X = 0$ or $Z_X^C = 1$. To keep the analysis here as simple as possible, we assume that discovery in carriage *only* impacts the pathogen's transmissibility from carriage, with no impact on the likelihood of subsequent infection or the pathogen's transmissibility during infection. In particular, there are no treatments available to clear the pathogen from carriage or to prevent progression to infection and there are no synergies between carriage HTC and infection HTC.

**Reproduction number.** Upon colonizing a host, strain $I_X$ of Pathogen 1 remains in an "undiscovered carriage phase" until discovery (rate $r_X$), progression to infection (rate $d^C$), or clearance (rate $\gamma^C + \sum_{x \notin X} \phi_x^C$). The undiscovered carriage phase therefore ends at rate $r_X + d^C + \gamma^C + \sum_{x \notin X} \phi_x^C$ and lasts on average for length of time $L_X^{C:U} = \frac{1}{r_X + d^C + \gamma^C + \sum_{x \notin X} \phi_x^C}$, during which transmission occurs at the uncontrolled rate $\beta^C$. The likelihood that strain $I_X$ is discovered in carriage is $\psi_X = \frac{r_X}{r_X + d^C + \gamma^C + \sum_{x \notin X} \phi_x^C}$. Upon discovery, the pathogen enters a "discovered carriage phase" that lasts until clearance or progression to infection. The discovered carriage phase, when it occurs, ends at rate $d^C + \gamma^C + \sum_{x \notin X} \phi_x^C$ and lasts on average for length of time $L_X^{C:D} = \frac{1}{d^C + \gamma^C + \sum_{x \notin X} \phi_x^C}$, during which transmission occurs at the lower controlled rate $Z_X^C \beta^C$. (Note that $L_X^{C:U} + \psi_X L_X^{C:D} = L_X^C$ as defined in Eq. C.3, reflecting our simplifying assumption that discovery does not speed clearance from carriage or slow progression to infection.) Strain $I_X$'s ex ante likelihood of progressing to cause infection $P_X^{C \to I} = \frac{d^C}{d^C + \gamma^C + \sum_{x \notin X} \phi_x^C}$, the same as in the main analysis. Overall, then, strain $I_X$'s reproduction number takes the form

$$R_{0,I_X} = \beta^C L_X^{C:U} + Z_X^C \beta^C \psi_X L_X^{C:D} + P_X^{C \to I} \Pi_X^I$$

*Equation C.6*

The first term in Eq. C.6 captures (ex ante expected) transmission during the undiscovered carriage phase, the second term captures transmission during the discovered carriage phase, and the last term captures transmission during infection, with $\Pi_X^I$ denoting the expected number of transmission events during infection (same as in Eq. C.1). Alternatively, $R_{0,X}$ can be equivalently expressed as

$$R_{0,X} = \left(\beta^C L_X^C + P_X^{C\to I}\Pi_X^I\right) - \psi_X\left(1 - Z_X^C\right)\beta^C L_X^{C:D}$$

*Equation C.7*

The first term in Eq. C.7 is strain $I_X$'s reproduction number when carriage RD is not available, the same as in Eq. C.1, while the second term is the amount by which carriage RD-enabled interventions *reduce* strain $I_X$'s reproduction number. To see why, note that strain $I_X$ is discovered with probability $\psi_X$ and, when discovered, is subjected to carriage HTC for expected length of time $L_X^{C:D}$, during which the transmission rate is reduced from $\beta^C$ to $Z_X^C\beta^C$. Thus, the overall effect is to reduce strain $I_X$'s reproduction number by $\psi_X\left(1 - Z_X^C\right)\beta^C L_X^{C:D}$.

<u>*Sensitive strain*</u>: By assumption, strain $I_0$ is not targeted for carriage HTC when discovered; thus, $Z_X^C = 1$ and $R_{0,0} = \frac{\beta^C + d^C\beta^I/\gamma_1}{d^C+\gamma^C+\phi_1^C+\phi_2^C}$, the same as before.

<u>*Drug-1-resistant strain*</u>: Strain $I_1$ is discovered while in carriage with probability $\psi_1 = \frac{r_1}{r_1+d^C+\gamma^C+\phi_2^C}$ and, if discovered, subjected to carriage HTC for expected length of time $L_1^{C:D} = \frac{1}{d^C+\gamma^C+\phi_2^C}$. By Eq. C.7, strain $I_1$'s reproduction number is now $R_{0,1} = \frac{\beta^C + d^C Z_1^I \beta^I/\gamma_2}{d^C+\gamma^C+\phi_2^C} - \frac{(1-Z_1^C)\beta^C r_1}{(r_1+d^C+\gamma^C+\phi_2^C)(d^C+\gamma^C+\phi_2^C)}$.

<u>*Drug-2-resistant strain*</u>: Strain $I_2$ is discovered with probability $\psi_2 = \frac{r_2}{r_2+d^C+\gamma^C+\phi_1^C}$ and, if discovered, subjected to carriage HTC for expected length of time $L_2^{C:D} = \frac{1}{d^C+\gamma^C+\phi_1^C}$. By Eq. C.7, strain $I_2$'s reproduction number is now $R_{0,2} = \frac{\beta^C + d^C Z_2^I \beta^I/\gamma_1}{d^C+\gamma^C+\phi_1^C} - \frac{(1-Z_2^C)\beta^C r_2}{(r_2+d^C+\gamma^C+\phi_1^C)(d^C+\gamma^C+\phi_1^C)}$.

<u>*Totally-resistant strain*</u>: Strain $I_{12}$ is discovered with probability $\psi_{12} = \frac{r_{12}}{r_{12}+d^C+\gamma^C}$ and, if discovered, subjected to carriage HTC for expected length of time $L_{12}^{C:D} = \frac{1}{d^C+\gamma^C}$. By Eq. C.7, strain $I_{12}$'s reproduction number is now $R_{0,12} = \frac{\beta^C + d^C Z_{12}^I \beta^I/\gamma_0}{d^C+\gamma^C} - \frac{(1-Z_{12}^C)\beta^C r_{12}}{(r_{12}+d^C+\gamma^C)(d^C+\gamma^C)}$.

### Part D. Inflows of drug-1-resistant infection

This section extends the obligate-pathogen model of Part B to allow for other mechanisms by which drug-1-resistant bacteria may enter the host population, namely, resistance-conferring mutation, competitive release, migration, and/or environmental infection. For analytical simplicity, we focus on the case in which (i) resistance to drug 2 has not yet emerged, so that only the sensitive strain $I_0$ and the drug-1-resistant strain $I_1$ are in circulation, (ii) there are no fitness costs associated with drug-1 resistance ($\beta_1 = \beta_0 = \beta$), (iii) drug 2 is equally effective as drug 1 ($\gamma_2 = \gamma_1$), and (iv) all infected hosts immediately come under medical attention.

The analysis in Part B established two key results in the baseline case without resistant inflows:

1. *Limited use of the second-line antibiotic:* Drug 2 can be sustainably reserved only for those with drug-1-resistant infection.
2. *Drug-1 resistant infection becomes / stays rare without needing to withhold treatment:* The prevalence of drug-1-resistant infection declines over time, so that drug-1 resistance becomes rare and then stays rare, even as all drug-1-sensitive infections are treated with drug 1.

The goal of this section is to determine whether these key findings continue to hold when there are inflows of drug-1-resistant infection (or simply "resistant infection"[7]) due to mutation, competitive release, migration, and/or environmental infection. Our answer, in essence, is that "it depends." When the inflow of resistant infection is small, we find that the long-run burden of resistant infection is small and hence that almost all patients can be sustainably treated with drug 1, much as in the baseline model without resistant inflows. On the other hand, very large inflows of resistant infection can be overwhelming, so that the long-run burden of drug-1-sensitive infection is small. In that case, doctors must eventually rely on drug 2 to treat almost all patients.

Although we focus for simplicity on a special case that abstracts from much of the complexity present in Part B, qualitatively similar results can be established through analogous extensions in which (i) drug-2 resistance and pan-resistance has emerged, (ii) drug 2 is less effective than drug 1, and (iii) infected hosts may not immediately come under medical attention.

---

[7] We focus here on drug-1-resistant infections, but similar analysis shows that our conclusions about when it is possible to select against drug-2-resistant and pan-resistant strains are also robust to small inflows. In particular, if pan-resistant infection is already widespread, adding a small inflow of pan-resistant infection has a small impact on maximal sustainable antibiotic use $\alpha_{12}^*$ computed in Part B.

**Case #1: Spontaneous emergence of resistance (e.g., mutation or competitive release)**

This section considers a variation of Part B's obligate-pathogen model in which each sensitive infection spontaneously transforms ("mutates") into a drug-1-resistant infection at rate $\delta^R > 0$. Although we use the language of mutation, the analysis here also applies to other mechanisms (such as horizontal gene transfer and competitive release) by which drug-1-resistant infection can emerge within a host who initially had drug-1-sensitive infection.

**Epidemiological dynamics**. At any given time, each host is in one of three epidemiologically-relevant states: $S$ (uninfected and susceptible to infection); $I_0$ (sensitive infection); or $I_1$ (resistant infection). Let $S(t)$ denote the population-wide mass of hosts in state $S$ and let $I_0(t), I_1(t)$ be the population-wide mass in states $I_0, I_1$ at time $t \geq 0$. Note that $S(t) + I_0(t) + I_1(t) = 1$ since the overall host population has mass one.

*Dynamics of sensitive infection.* Each sensitive infection transmits sensitive bacteria to a new host at rate $\beta$, fraction $S(t)$ of which are susceptible to infection. The flow of new sensitive infections is therefore $\beta I_0(t)S(t)$. So long as all sensitive infections are being treated, such infections clear at rate $\gamma_1$ (returning the host to state $S$) and mutate to become resistant at rate $\delta^R$ (transitioning to state $I_1$). The flow of hosts *out* of the sensitive-infected state $I_0$ is therefore $(\gamma_0 + \delta^R)I_0(t)$. Overall, then, the dynamics of $I_0(t)$ are characterized by the differential equation

$$\dot{I}_0(t) = \beta I_0(t)S(t) - (\gamma_1 + \delta^R)I_0(t)$$

*Equation D.1*

which can be expressed even more compactly in percentage-change terms:

$$\frac{\dot{I}_0(t)}{I_0(t)} = \beta S(t) - \gamma_1 - \delta^R$$

*Equation D.2*

*Dynamics of drug-1-resistant infection.* The dynamics of $I_1(t)$ are similarly derived, the only differences being: resistant infections have transmission rate $\beta Z$ as they are subjected to heightened transmission control (HTC); resistant infections are cleared at rate $\gamma_2$ since they are treated with drug 2; and the population-wide flow of sensitive infections that mutate to become resistant is $\delta^R I_0(t)$. All together,

$$\dot{I}_1(t) = \beta Z I_1(t)S(t) - \gamma_2 I_1(t) + \delta^R I_0(t)$$

*Equation D.3*

which in percentage-change terms takes the form

$$\frac{\dot{I}_1(t)}{I_1(t)} = \beta Z S(t) - \gamma_2 + \delta^R \frac{I_0(t)}{I_1(t)}$$

*Equation D.4*

**Long-run steady state**. Let $(S^*, I_0^*, I_1^*) = \lim_{t\to\infty}\big(S(t), I_0(t), I_1(t)\big)$ denote the long-run mass of hosts in each epidemiological state, also referred to as "the steady state."[8] An initial (purely mathematical) observation is that

$$\lim_{t\to\infty} \frac{\dot{I}_0(t)}{I_0(t)} = \beta S^* - \gamma_1 - \delta^R \leq 0$$

*Equation D.5*

$$\lim_{t\to\infty} \frac{\dot{I}_1(t)}{I_1(t)} = \beta Z S^* - \gamma_2 + \delta^R \frac{I_0^*}{I_1^*} \leq 0$$

*Equation D.6*

Moreover, it must be that $\lim_{t\to\infty} \frac{\dot{I}_0(t)}{I_0(t)} = 0$ if $I_0^* > 0$ and $\lim_{t\to\infty} \frac{\dot{I}_1(t)}{I_1(t)} = 0$ if $I_1^* > 0$. Because of the flow of new resistant infections due to mutation, it must be that $I_1^* > 0$; by Eq. D.6, we conclude that $S^* = \left(\gamma_2 - \delta^R \frac{I_0^*}{I_1^*}\right)/\beta Z$ and hence that $\lim_{t\to\infty} \frac{\dot{I}_0(t)}{I_0(t)} = \gamma_2/Z - \gamma_1 - \delta^R \left(1 + \frac{I_0^*}{I_1^*}/Z\right)$. Two relevant cases emerge.

*__When the sensitive strain is driven extinct ($I_0^* = 0$)__*. Suppose that $\delta^R > \bar{\delta}^R = \gamma_2/Z - \gamma_1$. Then $\lim_{t\to\infty} \frac{\dot{I}_0(t)}{I_0(t)} < 0$ and the burden of the sensitive infection must go to zero in the long run, despite the fact that strain $I_0$ has a strict reproductive advantage over strain $I_1$, as $R_{0,0} = \frac{\beta}{\gamma_1} > \frac{\beta Z}{\gamma_2} = R_{0,1}$.

*__When the sensitive strain is not driven extinct ($I_0^* > 0$)__*. Now, suppose that $\delta^R < \bar{\delta}^R$. In this case, if $I_0^* = 0$, then $\lim_{t\to\infty} \frac{\dot{I}_0(t)}{I_0(t)} = Z\gamma_2 - \gamma_1 - \delta^R > 0$, a contradiction; so, we conclude that $I_0^* > 0$ and hence that $\lim_{t\to\infty} \frac{\dot{I}_0(t)}{I_0(t)} = 0$. By Eq. D.5, we conclude that

[8] One can show that the limit defining the steady-state exists and does not depend on initial conditions. Mathematical details to establish these points (here and in the more complicated case analyzed later) are straightforward and omitted to save space.

$$S^* = \frac{\gamma_1 + \delta^R}{\beta}$$

*Equation D.7*

and hence that the long-run burden of infection is $I_0^* + I_1^* = 1 - S^* = \frac{\beta - \gamma_1 - \delta^R}{\beta}$. Moreover, using the fact that $S^*$ also equals $\left(\gamma_2 - \delta^R \frac{I_0^*}{I_1^*}\right)/\beta Z$, we conclude that

$$\frac{I_1^*}{I_0^*} = \frac{\delta^R}{\gamma_2 - (\gamma_1 + \delta^R)Z}$$

*Equation D.8*

_Discussion: Impact of rare mutation on long-run outcomes._ For small mutation rates $\delta^R \approx 0$, the long-run burden of drug-1-sensitive infection $I_0^* \approx \frac{\beta - \gamma_1}{\beta}$, the same as in a benchmark model without mutation, while the long-run burden of drug-1-resistant infection is approximately proportional to $\delta^R$, with $\frac{I_1^*}{\delta^R} \approx \frac{\beta - \gamma_1}{\beta(\gamma_2 - (\gamma_1 + \delta^R)Z)}$ . In particular, so long as resistance-conferring mutation is rare, resistant infection will also be rare.

**Case #2: Migration of hosts with resistant infection**

This section considers a different extension of Part B's obligate-pathogen model in which a constant flow of $\delta^M > 0$ "migrants" enter the host population with drug-1-resistant infection while, to keep the population-size fixed, each host exits at the same rate $\delta^M$. The dynamics of sensitive and drug-1-resistant infection are now characterized by differential equations $\dot{I}_0(t) = \beta I_0(t)S(t) - (\gamma_1 + \delta^M)I_0(t)$ and $\dot{I}_1(t) = \beta Z I_1(t)S(t) - (\gamma_2 + \delta^R)I_1(t) + \delta^R$ which in percentage-change terms can be expressed as

$$\frac{\dot{I}_0(t)}{I_0(t)} = \beta S(t) - \gamma_1 - \delta^M$$

*Equation D.9*

$$\frac{\dot{I}_1(t)}{I_1(t)} = \beta Z S(t) - \gamma_2 - \delta^M + \frac{\delta^M}{I_1(t)}$$

*Equation D.10*

Because of the flow of new resistant infections due to migration, it must be that $I_1^* > 0$; by Eq. D.10, we conclude that $S^* = \left(\gamma_2 - \delta^M \frac{1-I_1^*}{I_1^*}\right)/\beta Z$ and hence $\lim_{t\to\infty} \frac{\dot{I}_0(t)}{I_0(t)} = \gamma_2/Z - \gamma_1 - \delta^M \left(1 + \frac{1-I_1^*}{I_1^*}/Z\right)$. Two relevant cases emerge.

*When the sensitive strain is driven extinct ($I_0^* = 0$).* Suppose that $\delta^M > \bar{\delta}^M = \gamma_2/Z - \gamma_1$, the same critical level as in the earlier mutation analysis. Then $\lim_{t\to\infty} \frac{\dot{I}_0(t)}{I_0(t)} < 0$ and the burden of multi-drug-sensitive infection must go to zero in the long run.

*When the sensitive strain is not driven extinct ($I_0^* > 0$).* Now, suppose that $\delta^M < \bar{\delta}^M$ If $I_0^* = 0$, then $\lim_{t\to\infty} \frac{\dot{I}_0(t)}{I_0(t)} = \gamma_2/Z - \gamma_1 - \delta^R > 0$, a contradiction; so, we conclude that $I_0^* > 0$ and hence that $\lim_{t\to\infty} \frac{\dot{I}_0(t)}{I_0(t)} = 0$. By Eq. D.9, we conclude that

$$S^* = \frac{\gamma_1 + \delta^M}{\beta}$$

*Equation D.11*

and hence that $I_0^* + I_1^* = \frac{\beta - \gamma_1 - \delta^R}{\beta}$. Moreover, using the fact that $S^*$ also equals $\left(\gamma_2 - \delta^M \frac{1-I_1^*}{I_1^*}\right)/\beta Z$, we conclude that

$$\frac{I_1^*}{1 - I_1^*} = \frac{\delta^M}{\gamma_2 - (\gamma_1 + \delta^M)Z}$$

*Equation D.12*

*Discussion: mutation vs migration.* Suppose that the resistance-conferring mutation rate in Case #1 is equal to the in-migration rate of resistant hosts here, i.e., $\delta^R = \delta^M = \delta$, and let $(S^{*R}, I_0^{*R}, I_1^{*R})$ and $(S^{*M}, I_0^{*M}, I_1^{*M})$ denote the steady state in each case. The previous analysis implies that (i) the long-run burden of infection is the same whether resistant infection enters the population through mutation or migration but (ii) the long-run prevalence of resistance is higher in the case of migration. Why? First, by Eqs. D.7 and D.11, $I_0^{*R} + I_1^{*R} = I_0^{*M} + I_1^{*M} = 1 - \frac{\gamma_1 + \delta}{\beta}$; so, the long-run burden of infection is the same. Next, by Eqs. D.8 and D.12, $\frac{1-I_1^{*M}}{I_1^{*M}} = \frac{I_0^{*R}}{I_1^{*R}}$ and hence $\frac{1}{I_1^{*M}} = 1 + \frac{1-I_1^{*M}}{I_1^{*M}} = 1 + \frac{I_0^{*R}}{I_1^{*R}} = \frac{1-S^{*R}}{I_1^{*R}}$, implying that $I_1^{*R} = (1 - S^{*R})I_1^{*M} < I_1^{*M}$; so, the long-run burden of resistant infection is higher under migration.

#### Case #3: Resistant environmental infection

This section considers yet another extension of the obligate-pathogen model, in which susceptible hosts acquire drug-1-resistant infection from environmental sources at constant rate $\delta^E > 0$. The dynamics of sensitive and drug-1-resistant infection are now characterized by differential equations $\dot{I}_0(t) = \beta I_0(t)S(t) - \gamma_1 I_0(t)$ and $\dot{I}_1(t) = \beta Z I_1(t)S(t) - \gamma_2 I_1(t) + \delta^E S(t)$, so that

$$\frac{\dot{I}_0(t)}{I_0(t)} = \beta S(t) - \gamma_1$$

*Equation D.13*

$$\frac{\dot{I}_1(t)}{I_1(t)} = \beta Z S(t) - \gamma_2 + \delta^E \frac{S(t)}{I_1(t)}$$

*Equation D.14*

Because of the flow of new resistant infections due to environmental infection, it must be that $I_1^* > 0$ and hence that the steady-state condition $\lim_{t\to\infty} \frac{\dot{I}_1(t)}{I_1(t)} = 0$ holds. By Eq. D.14, this implies

$$\frac{I_1^*}{S^*} = \frac{\delta^E}{\gamma_2 - \beta Z S^*}$$

*Equation D.15*

There is a threshold $\bar{\delta}^E > 0$ such that $I_0^* > 0$ when $\delta^E < \bar{\delta}^E$ but $I_0^* = 0$ when $\delta^E \geq \bar{\delta}^E$. To derive $\bar{\delta}^E$, suppose for the moment that $I_0^* > 0$, so that the steady-state condition $\lim_{t\to\infty} \frac{\dot{I}_0(t)}{I_0(t)} = 0$ must hold. By Eq. D.13, this implies $S^* = \frac{\gamma_1}{\beta}$; by Eq. D.15, this is turn implies $I_1^* = \delta^E \frac{\gamma_1}{\beta\left(\gamma_2 - \frac{\gamma_1}{Z}\right)}$.

Since the host population has mass one, $I_0^* > 0$ also implies $S^* + I_1^* < 1$. However, this is only true when $\delta^E < \bar{\delta}^E = \frac{\beta - \gamma_1}{\gamma_1}(\gamma_2 - Z\gamma_1)$. We are left with two logical possibilities. *Case #1:* $\delta^E < \bar{\delta}^E$ *and* $I_0^* > 0$. The steady state in this case is as previously derived:

$$S^* = \frac{\gamma_1}{\beta} \text{ and } I_1^* = \delta^E \frac{\gamma_1}{\beta(\gamma_2 - Z\gamma_1)}$$

*Equation D.16*

and $I_0^* = 1 - S^* - I_1^*$. *Case #2:* $\delta^E \geq \bar{\delta}^E$ *and* $I_0^* = 0$. In this case, $I_1^* = 1 - S^*$ and, by Eq. D.14, the steady-state condition $\lim_{t\to\infty} \frac{\dot{I}_1(t)}{I_1(t)} = 0$ can be written as

$$\lim_{t\to\infty} \frac{\dot{I}_1(t)}{I_1(t)} = \beta Z S^* - \gamma_2 + \delta^E \frac{S^*}{1 - S^*} = 0$$

*Equation D.17*

which uniquely characterizes $S^*$. (Mathematical details to show that Eq. D.17 has a unique solution in $S^*$ are straightforward and omitted.)

Discussion: environmental infection vs migration. Suppose that the environmental infection rate here is equal to the in-migration rate of resistant hosts in Case #2 and low enough that the sensitive strain is not driven extinct, i.e., $\delta^E = \delta^M = \delta < \min\{\bar{\delta}^M, \bar{\delta}^E\}$ and let $(S^{*E}, I_0^{*E}, I_1^{*E})$ and $(S^{*M}, I_0^{*M}, I_1^{*M})$ denote the steady state in each case. Our analysis implies that (i) the long-run burden of infection is higher when resistance enters the population through environmental infection but (ii) the long-run burden of resistant infection is higher when resistance enters the population through migration. Why? First, recall that $S^{*E} = \frac{\gamma_1}{\beta} < \frac{\gamma_1+\delta}{\beta} = S^{*M}$; so, $I_0^{*E} + I_1^{*E} = 1 - \frac{\gamma_1}{\beta} > 1 - \frac{\gamma_1+\delta}{\beta} = I_0^{*M} + I_1^{*M}$. Next, by Eqs. D.12 and D.15, $\frac{I_1^{*M}}{1-I_1^{*M}} = \frac{\delta}{\gamma_2-(\gamma_1+\delta)Z} > \frac{\delta}{\gamma_2-\gamma_1 Z} = \frac{I^{*E}}{S^{*E}}$. Since $S^{*E} < S^{*M} < 1 - I_1^{*M}$, we conclude $I_1^{*M} > I_1^{*E}$.

### Part E. Extension: diagnostic failures

This section extends the obligate-pathogen model to allow for the possibility of "diagnostic error" (that POC-RD may sometimes return incorrect results) and "diagnostic escape" (that new bacterial strains may emerge that cannot be detected until POC-RD testing is updated to identify them).

#### Case #1: Diagnostic error

Any resistance diagnostic will be subject to occasional error. In particular, suppose that infections that are resistant to drug 1 will be wrongly labeled by POC-RD as "sensitive" with probability $\delta^{I}$ ("type-I error") while those that are sensitive to drug 1 will be wrongly labeled "resistant" with probability $\delta^{II}$ ("type-II error"), and similarly for drug-2 resistance. So, for instance, an infection that is sensitive to both antibiotics will be wrongly labeled as "pan-resistant" whenever two type-II errors happen at the same time, i.e., with probability $(\delta^{II})^2$. Some resistance diagnostics frequently return erroneous results; see e.g. (1) on error in culture-based CRE diagnosis. However, recently-developed molecular diagnostics can detect the presence of resistance-conferring genes with very little error; see e.g., (2) and (3) on the Xpert MRSA test for detecting methicillin-resistant staph and (4) on the Xpert MRDO assay for detecting carbapenase-producing gram-negative bacilli. Bearing this in mind, we focus here on the case in which diagnostic error is rare, i.e., $\delta^{I} \approx 0$ and $\delta^{II} \approx 0$.

Diagnostic error has two main sorts of effects. First, diagnostic error dilutes the effectiveness of resistance-targeted medical intervention, since some sensitive infections are controlled more intensively and some resistant infections less intensively than if POC-RD were error-free. However, so long as the error rates $\delta^{I}$, $\delta^{II}$ are small, each strain's reproduction number will be approximately the same as if there were no diagnostic error. In particular, any resistance-targeted intervention that puts the pan-sensitive strain at a strict reproductive advantage absent diagnostic error will continue to do so when diagnostic error is rare.

Second, and perhaps more important in practice, diagnostic error can undermine doctors' confidence in the results of POC-RD testing, especially when resistant infection is even rarer than diagnostic error. To see the point, suppose that the burden of each resistant strain has been successfully reduced so much so that $\frac{I_1(t)}{\delta^{II}}, \frac{I_2(t)}{\delta^{II}}, \frac{I_{12}(t)}{(\delta^{II})^2} \approx 0$, and that an infection is then identified by POC-RD as "pan-resistant." This test result could be true (if the infecting strain is $I_{12}$) or could be due to a single type-II error (if the infecting strain is $I_1$ or $I_2$) or two type-II errors (if the infecting strain is $I_0$). Applying Bayes' Rule, the conditional probability that an infection found to be "pan-resistant" really is pan-resistant is

$$\Pr(I_{12}|\text{"pan resistant"}) = \frac{(1-\delta^{II})^2 I_{12}(t)}{(1-\delta^{II})^2 I_{12}(t) + \delta^{II}(1-\delta^{I})\big(I_1(t)+I_2(t)\big) + (\delta^{II})^2 I_0(t)} \approx 0$$

*Equation E.1*

leaving the doctor with essentially the same (lack of) information as if the test had not been conducted.

In this context, policy of isolating all patients found to be "pan-resistant" would put the pan-resistant strain at a reproductive disadvantage relative to the other strains, ensuring that pan-resistant infection remains rare. (Why? Fraction $(1-\delta^{I})^2 \approx 100\%$ of sensitive infections would be isolated under such a policy, while only fraction $\delta^{II}(1-\delta^{I}) \approx 0$ of one-drug-resistant infections and fraction $(\delta^{II})^2 \approx 0$ of sensitive infections would be isolated.) Even so, doctors may naturally hesitate to order intensive interventions such as isolation unless they are sufficiently confident that the patient's infection actually is pan-resistant. If so, totally-resistant infections will not be effectively controlled until they have become sufficiently common that $\Pr(I_{12}|\text{"totally resistant"})$ exceeds the threshold for isolation. For instance, suppose that doctors need 90% certainty that an infection is caused by strain $I_{12}$ before they are willing to isolate a patient, that type-I and type-II errors each occur 10% of the time, and that both one-drug-resistant strains are exceedingly rare. Eq. E.1 then becomes $\Pr\big(I_{12}|\text{"pan-resistant"}\big) = \frac{(0.9)^2 I_{12}(t)}{(0.9)^2 I_{12}(t)+(0.1)^2 I_0(t)}$. This exceeds 90% only when $\frac{I_{12}(t)}{I_0(t)} > \frac{1}{9}$, i.e. when the pan-resistant strain is already sufficiently widespread to be causing 10% of all infections.

**Case #2: Diagnostic escape**

Resistant bacteria may occasionally mutate in ways that allow them to maintain their antibiotic resistance but evade detection by current POC-RD; see e.g. (5) on the emergence of MRSA strains that are not detectable by the Xpert MRSA test due to recombinations within the staphylococcal cassette chromosome *mec*. During the period of time before a new resistant strain becomes detectable (by being identified through population-wide surveillance and added to the set of strains that can be found by POC-RD), it will enjoy a reproductive advantage and the burden of infection it causes will increase over time. However, once that initial period has passed and the new strain has become detectable, its burden of infection will evolve over time much like any other resistant strain. In particular, if the conditions are in place to reverse the rise of detectable resistance—as discussed in Part B for obligate pathogens and Part C for opportunistic pathogens—the prevalence of the new strain will then decline over time until it eventually becomes a negligible source of infection. Our results are therefore robust to the possibility of diagnostic escape, so long as new resistant strains can be quickly identified and POC-RD can be quickly updated to detect them.

To flesh out this point, consider an especially simple scenario with two equally-effective antibiotics ($\gamma_2 = \gamma_1 > \gamma_0$) and no fitness costs ($\beta_1 = \beta_0 = \beta$) in which resistance to drug 2 has not yet emerged, POC-RD is available that can detect all drug-1-resistant strains currently in circulation, and heightened measures are available to reduce transmission from infections found to be drug-1 resistant ($Z > 1$). As shown in Part B, POC-RD enables all detectable drug-1-resistant strains to be held at a strict reproductive disadvantage, as sensitive strains are treated with drug 1 while drug-1-resistant strains are treated with drug 2 and subjected to HTC.

*Starting point: close to the steady state without resistance*. Let $S(t)$ be the mass of hosts that are uninfected, $I_0(t)$ the mass of hosts infected with a sensitive strain, and $I_1(t)$ the mass of hosts infected with a drug-1-resistant strain that is detectable by POC-RD. Prior to diagnostic escape, $S(t) + I_0(t) + I_1(t) = 1$ and epidemiological dynamics are determined by the following system of differential equations

$$\frac{\dot{I}_0(t)}{I_0(t)} = \beta S(t) - \gamma_1$$

*Equation E.2*

$$\frac{\dot{I}_1(t)}{I_1(t)} = \beta Z S(t) - \gamma_2$$

*Equation E.3*

In Part D, we derived the long-run steady state of this dynamic system: $\lim_{t\to\infty} S(t) = S^* = \frac{\gamma_1}{\beta}$, $\lim_{t\to\infty} I_0(t) = I_0^* = 1 - \frac{\gamma_1}{\beta}$, and $\lim_{t\to\infty} I_1(t) = 0$. Suppose that this steady state has been approximately reached, when a new drug-1-resistant strain emerges that cannot be detected by POC-RD. (Our focus here is on diagnostic escape, but the analysis equally applies to settings in which unknown genetic determinants of drug-1 resistance enter the disease-causing bacterial population through horizontal gene transfer, host migration, or other means.)

*Exponential increase of newly-emerged resistant strain*. Let $I_1^{YES}(t)$ denote the burden of infection caused by (preexisting) detectable drug-1-resistant strains and let $I_1^{NO}(t)$ denote the burden caused by a (new) undetectable drug-1-resistant strain. As a normalization, define "time 0" as the moment that the undetectable drug-1-resistant strain has grown sufficiently prevalent to be infecting mass $\Delta \approx 0$ of hosts. So, $S(0) \approx \frac{\gamma_1}{\beta}$, $I_0(0) \approx 1 - \frac{\gamma_1}{\beta}$, $I_1^{YES}(0) \approx 0$, and $I_1^{NO}(0) = \Delta$. Epidemiological dynamics of $I_0(t), I_1^{YES}(t)$ are still determined by Eqs. E.2 and E.3, while the dynamics of undetectable drug-1-resistant infection are determined by

$$\frac{\dot{I}_1^{NO}(t)}{I_1^{NO}(t)} = \beta S(t) - \gamma_0 \approx \gamma_1 - \gamma_0$$

*Equation E.4*

By Eq. E.4, the burden of infection caused by the newly-emerged drug-1-resistant strain increases approximately exponentially at rate $\gamma_1 - \gamma_0$;[9] so, at each time $t > 0$, $I_1^{NO}(t) \approx \Delta e^{t(\gamma_1 - \gamma_0)}$ and the *total burden* of the newly-emerged strain from time 0 until time $t$ equals

$$\text{Total Burden during time-period } [0, t] \approx \Delta \int_0^t e^{\hat{t}(\gamma_1 - \gamma_0)} d\hat{t} = \frac{\Delta}{\gamma_1 - \gamma_0}\left(e^{t(\gamma_1 - \gamma_0)} - 1\right)$$

*Equation E.5*

*Exponential decrease of newly-detectable resistant strain*. Suppose that it takes time $T > 0$ for public-health officials to identify a newly-emerged drug-1-resistant strain through population-wide resistance surveillance and for diagnostic-makers to update POC-RD so that the new strain can be detected at the point of care.[10] As shown previously, $I_1^{NO}(T) \approx \Delta e^{T(\gamma_1 - \gamma_0)}$. Starting at time $T$, infections caused by the new drug-1-resistant strain will now be treated with drug 2 and subjected to HTC, and hence be subject to the same epidemiological dynamics as all of the other detectable drug-1-resistant strain, i.e., $\frac{\dot{I}_1^{YES}(t)}{I_1^{YES}(t)} \approx \beta Z S^* - \gamma_2 = Z\gamma_1 - \gamma_2$ (same as Eq. E.3). Since both antibiotics are by assumption equally

[9] In this section, we make extensive use of a basic mathematical fact, that only the exponential function has a constant percentage-rate of change. That is, if $\frac{df(t)/dt}{f(t)} = C$ for all $t$, then it must be that $f(t) = f(0)e^{tC}$, where $C$ is the rate of exponential increase (or, if $C < 0$, $-C$ is the rate of exponential decrease).

[10] The manner of diagnostic escape and of diagnostic updating depends on the nature of the test. For simplicity, consider a setting in which POC-RD works by rapidly sequencing the genome of the infecting bacteria and then comparing it against a database of known resistant strains (or known genetic determinants of resistance). POC-RD is "updated" in this context by simply adding the sequence of a newly-identified resistant strain to a database of known resistant strains. Such whole-genome sequence databases already exist, e.g. the WGS NARMS database of known resistant strains of enteric bacteria (6), but point-of-care diagnostics that link directly to such databases are not yet available.

effective, $Z\gamma_1 - \gamma_2 < 0$ and the burden of infection caused by the new strain declines approximately exponentially over time at rate $\gamma_2 - Z\gamma_1$. In particular, $I_1^{YES}(t) \approx I_1^{YES}(T) \times e^{-(t-T)(\gamma_2 - Z\gamma_1)}$ at each time $t > T$. Given this exponential decline, the subsequent burden of infection caused by the new resistant strain is

$$\text{Total Burden during time-period } [T, \infty) \approx I_1^{YES}(T) \int_T^{\infty} e^{-(t-T)(\gamma_2 - Z\gamma_1)} dt = \frac{\Delta e^{T(\gamma_1 - \gamma_0)}}{\gamma_2 - Z\gamma_1}$$

*Equation E.6*

Combining Eqs. E.5 and E.6, note that the newly-emerged drug-1-resistant imposes a *finite* total burden on the host population over time:

$$\text{Total Burden during time-period } [0, \infty) = \Delta \left( \frac{e^{T(\gamma_1 - \gamma_0)} - 1}{\gamma_1 - \gamma_0} + \frac{e^{T(\gamma_1 - \gamma_0)}}{\gamma_2 - Z\gamma_1} \right)$$

*Equation E.7*

By contrast, the burden of the sensitive strain during any time-period $[0, t]$ is approximately $tS^* > 0$, dwarfing the impact of each "diagnostic escapee."

nalAntimicrobialResistanceMonitoringSystem/UCM528861.pdf.

### Part F. Basis for Fig 3

Methods and data sources underlying point estimates and confidence intervals for all organisms except *C. difficile* are described in Tedijanto, et al. (2018). Confidence intervals were not calculated for *N. gonorrhoeae* and *C. difficile*, which required different data sources and methodology from all other organisms due to the rarity of disease. All estimates are based on ambulatory care prescriptions and diagnoses from 2010-2011 in the United States, except for *C. difficile*, for which details are described below. US census estimates for 2010 and 2011 were used to calculate rates per person-year.

***C. difficile*** **methodology and data sources**

Target antibiotic exposures

The number of incident cases of *C. difficile* infection in 2011 was estimated to be 453,000 (Lessa et al. 2015). We assumed the same number of incident cases occurred in 2010 and that each case was associated with one target antibiotic "exposure" (one prescription, in the ambulatory care setting).

Bystander antibiotic exposures

Based on a recent review, which found the prevalence of asymptomatic *C. difficile* colonization rates among healthy adults in the general population to be between 0 and 15% (Furuya-Kanamori, et al. 2015), we estimated carriage prevalence to be 7.5%. The rate of bystander antibiotic exposures was calculated as the product of carriage prevalence and the rate of total antibiotic exposures in US ambulatory care, based on US census estimates and the National Ambulatory Medical Care Survey and National Hospital Ambulatory Medical Care Survey (NAMCS/NHAMCS) from 2010-11.

Furuya-Kanamori, L., Marquess, J., Yakob, L., Riley, T. V., Paterson, D. L., Foster, N. F., ... & Clements, A.C. (2015). Asymptomatic Clostridium difficile colonization: epidemiology and clinical implications. *BMC infectious diseases*, *15*(1), 516.

Lessa, F. C., Mu, Y., Bamberg, W. M., Beldavs, Z. G., Dumyati, G. K., Dunn, J. R., ... & Wilson, L. E. (2015). Burden of Clostridium difficile infection in the United States. *New England Journal of Medicine*, *372*(9), 825-834.

Tedijanto, C., Olesen, S., Grad, Y., & Lipsitch, M. (2018). Estimating the proportion of bystander selection for antibiotic resistance in the US. PNAS, 115(51), E11988-E11995.

United States, U.S. Census Bureau. (2011). *Statistical abstract of the United States, 2011.* Washington, DC.

### Part G. Basis for pneumococcal parameter values in Figs 4,5

**For parameter definitions, see Table 1 in the main text. All rates are defined per day.**

***d*=0.001**

The disease-development rate (*d*) is calculated as the incidence of disease over the prevalence of carriage, that is, (76 infections / 1000 child years) / (200 carriages / 1000 children) = 0.38 infections per year, that is, 0.001 infections per day. The bases for these incidence and prevalence values are given below.

Incidence of disease. Disease incidence data are from a nationwide study of ambulatory care in the USA (*1*). The age group used is individuals 0-19 years old.

*Sinusitis*. There were 65 (CI95 51-79) antibiotic prescriptions for sinusitis per 1000 child years, and 85% of cases were prescribed antibiotics (*1*). The incidence of sinusitis is thus 76 cases per 1000 child years. According to a meta-analysis of causative agents of acute bacterial rhinosinusitis (not restricted to children) (*2*) 24% of cases were due to *S. pneumoniae* (calculated from table 2 in that paper as a weighted average across studies from the USA, for which the number of patients was given, with each study weighted by the number of patients included). The incidence of pneumococcal sinusitis is thus estimated as approximately *18 cases per 1000 child years*. Since (*2*) only included bacterial sinusitis (not viral), this number may be an overestimate.

*Suppurative otitis media*. There were 154 (CI95 131-177) antibiotic prescriptions per 1000 child years, and 82% of cases were prescribed antibiotics (*1*). The incidence of suppurative otitis media is thus 188 cases per 1000 child years. A systematic review of the causative agents of otitis media in children (*3*) found that *S. pneumoniae* was responsible for on average 29% (range 20.6-38.4) of cases in North America. The incidence of pneumococcal otitis media is thus approximately *55 cases per 1000 child years*.

*Pneumonia*. There were 22 (CI95 16-27) antibiotic prescriptions per 1000 child years, and 79% of cases were prescribed antibiotics (*1*). The incidence of pneumonia is thus 28 cases per 1000 child years. A study of pneumonia in children and adolescents found that 11% were due to *S. pneumoniae* (*4*). The incidence of pneumococcal pneumonia is thus approximately *3 cases per 1000 child years*. (It is plausible that the pneumonia cases that are treated are a non-random subset, and that up to 100% of pneumococcal cases are treated. However, since pneumonia is relatively rare, and a high proportion of pneumonia cases are treated, this would have little effect on the numbers calculated herein.)

The incidence of pneumococcal disease is calculated as the sum of the above incidences, that is, approximately 76 cases per 1000 child years.

Prevalence of carriage. In a meta-analysis (*5*) the median for studies in children less than 5 years old was 53%. For children 5-17 years old the median was 27%. (Calculated from Table 1 in that paper.) A study of UK families reported approximately 50% prevalence in preschool children (52% for 0-2 and 45% for children 4-5 years old) (*6*). A study of children attending one US hospital for other reasons reported 34% for children 0-6 years old, and 18% for children 0-17 years old (*7*). We conclude that for USA and similar countries a prevalence of carriage of approximately 400 per 1000 preschool children is reasonable. For children 0-17 years old the prevalence is lower. Since disease and prescription data (above) are for children 0-19 years old, we will use 200 cases per 1000 children. This is, of course, very approximate.

**$\phi_1^C$=0.0005**

The rate of bystander exposure to drug 1 for drug susceptible pneumococcus is taken from (*9*).

**$\phi_2^C$=0**

We make the assumption that bystander exposure to drug 2 is minimal, given a rarely used drug 2 option. In practice, values of $\phi_2^C$ in the region of $\phi_1^C$ have minimal quantitative impact on drug 1 resistance selection, as both drug 1 resistant and drug 1 susceptible strains are hit equally by any drug 2 exposures.

**$\gamma_0^I$=0.125**

Pneumococcal disease (in the USA) is dominated by otitis media and sinusitis, pneumonia making a minor contribution (above). The duration of culture positivity in acute otorrhea without antibiotic treatment was investigated by (*10*), and reported as a median of 8 days (interquartile range 2.25-8) (per protocol). However, this study was not limited to pneumococci, and the children were only followed for one week, 8 days duration being registered if the child was culture positive at the end of the period. The duration of sinusitis was investigated by (*11*), and was reported as a mean of 6.4 days. However, this referred only to symptoms, not culture positivity. Since the relevant duration for our model is the duration of transmissibility, which requires live bacteria, we use the result of culture positivity in otitis, that is, 8 days. A duration of 8 days yields a clearance rate of 1/8=0.125 episodes per day.

**$\gamma_1^I$=1**

The duration of culture positivity in acute otorrhea with antibiotic treatment (amoxicillin-clavulanate) was investigated by (*10*), and reported as a median of 1 day (interquartile range 1-2) (per protocol). However, this study was not limited to pneumococci. The duration of sinusitis was investigated by (*11*), and was reported as a mean of 6.0 days. However, this referred only to symptoms, not culture positivity. Since the relevant duration for our model is the duration of transmissibility, which requires live bacteria, we use the result of culture positivity in otitis, that is, 1 day. A duration of 1 day yields a clearance rate of 1/1=1 episode per day.

**Duration of carriage.**

The estimates of capsular specific duration of carriage were derived from *(12).* For serotypes 1,4 and 5 their median carriage duration was reported to be less than the sampling resolution of 4 weeks. We therefore estimate carriage of 2 weeks for these serotypes, as the midpoint of the 0-4 weeks uncertainty.
